# The Group B Streptococcal surface antigen I/II protein, BspC, interacts with host vimentin to promote adherence to brain endothelium and inflammation during the pathogenesis of meningitis

**DOI:** 10.1101/544395

**Authors:** Liwen Deng, Brady L. Spencer, Joshua A. Holmes, Rong Mu, Sara Rego, Thomas A. Weston, Yoonsung Hu, Glenda F. Sanches, Sunghyun Yoon, Nogi Park, Prescilla E. Nagao, Howard F. Jenkinson, Justin A. Thornton, Keun Seok Seo, Angela H. Nobbs, Kelly S. Doran

## Abstract

*Streptococcus agalactiae* (Group B *Streptococcus*, GBS) normally colonizes healthy adults but can cause invasive disease, such as meningitis, in the newborn. To gain access to the central nervous system, GBS must interact with and penetrate brain or meningeal blood vessels; however, the exact mechanisms are still being elucidated. Here, we investigate the contribution of BspC, an antigen I/II family adhesin, to the pathogenesis of GBS meningitis. Disruption of the *bspC* gene reduced GBS adherence to human cerebral microvascular endothelial cells (hCMEC), while heterologous expression of BspC in non-adherent *Lactococcus lactis* conferred bacterial attachment. In a murine model of hematogenous meningitis, mice infected with Δ*bspC* mutants exhibited lower mortality as well as decreased brain bacterial counts and inflammatory infiltrate compared with mice infected with WT GBS strains. Further, BspC was both necessary and sufficient to induce neutrophil chemokine expression. We determined that BspC interacts with the host cytoskeleton component vimentin, and confirmed this interaction using a bacterial two-hybrid assay, microscale thermophoresis, immunofluorescent staining, and imaging flow cytometry. Vimentin null mice were protected from WT GBS infection and also exhibited less inflammatory cytokine production in brain tissue. These results suggest that BspC and the vimentin interaction is critical for the pathogenesis of GBS meningitis.

**AUTHOR SUMMARY:** Group B *Streptococcus* (GBS) typically colonizes healthy adults but can cause severe disease in immune compromised individuals, including newborns. Despite wide-spread intrapartum antibiotic prophylaxis given to pregnant women, GBS remains a leading cause of neonatal meningitis. To cause meningitis, GBS must interact with and penetrate the blood-brain barrier (BBB), which separates bacteria and immune cells in the blood from the brain. In order to develop targeted therapies to treat GBS meningitis, it is important to understand the mechanisms of BBB crossing. Here, we describe the role of the GBS surface factor, BspC, in promoting meningitis and discover the host ligand for BspC, vimentin, which is an intermediate filament protein that is constitutively expressed by endothelial cells. We determined that BspC interacts with the C-terminal domain of cell-surface vimentin to promote bacterial attachment to brain endothelial cells and that purified BspC protein can induce immune signaling pathways. In a mouse model of hematogenous meningitis, we observed that a GBS mutant lacking BspC was less virulent compared to WT GBS and resulted in less inflammatory disease. We also observed that mice lacking vimentin were protected from GBS infection. These results reveal the importance of the BspC-vimentin interaction in the progression of GBS meningitis disease.

## INTRODUCTION

*Streptococcus agalactiae* (Group B *Streptococcus*, GBS) is an opportunistic pathogen that asymptomatically colonizes the vaginal tract of up to 30% of healthy women. However, GBS possesses a variety of virulence factors and can cause severe disease when transmitted to susceptible hosts such as the newborn. Despite widespread intrapartum antibiotic administration to colonized mothers, GBS remains a leading cause of pneumonia, sepsis, and meningitis in neonates (1, 2). Bacterial meningitis is a life-threatening infection of the central nervous system (CNS) and is marked by transit of the bacterium across endothelial barriers, such as the blood-brain barrier (BBB) or the meningeal blood-cerebral spinal fluid barrier (mBCSFB). Both consist of a single layer of specialized endothelial cells that serve to maintain brain homeostasis and generally prevent pathogen entry into the CNS (3–5). Symptoms of bacterial meningitis may be due to the combined effect of bacterial adherence and brain penetration, direct cellular injury caused by bacterial cytotoxins, and/or activation of host inflammatory pathways that can disrupt brain barrier integrity and damage underlying nervous tissue. (6–8)

Bacterial meningitis typically develops as a result of the pathogen spreading from the blood to the meninges. In order to disseminate from the blood into the brain, GBS must first interact with barrier endothelial cells (9). A number of surface-associated factors that contribute to GBS-brain endothelium interactions have been described such as lipoteichoic acid (LTA) (10), pili (11), serine-rich repeat proteins (Srr) (12), and streptococcal fibronectin-binding protein (SfbA) (13). Pili, the Srr proteins, and SfbA have been shown to interact with extracellular matrix (ECM) components, which may help to bridge to host receptors such as integrins or other ECM receptors. However, a direct interaction between a GBS adhesin and an endothelial cell receptor has not been described.

Antigen I/II family (AgI/II) proteins are multifunctional adhesins that have been well characterized as colonization determinants of oral streptococci (14). These proteins mediate attachment of *Streptococcus mutans* and *Streptococcus gordonii* to tooth surfaces and can stimulate an immune response from the colonized host (14). Genes encoding AgI/II polypeptides are found in streptococcal species indigenous to the human mouth as well as other pathogenic streptococci such as GBS, *S. pyogenes* (Group A *Streptococcus*, GAS), and *S. suis* (14, 15). Intriguingly, the GAS AgI/II protein AspA (Group <u>A</u> *Streptococcus* <u>s</u>urface <u>p</u>rotein) is absent in many GAS M serotypes and is found predominantly among M serotypes implicated in puerperal sepsis and neonatal infections, including M2, M4, and M28. The gene encoding AspA is located within an integrative and conjugative element designated region of difference 2 (RD2), which likely originated in GBS and was acquired by invasive GAS serotypes through horizontal gene transfer (Fig. 1A). It has been proposed that genes carried within RD2 may contribute to pathogenicity of both GAS and GBS in pregnant women and newborns (16, 17). Supporting this, AspA has been shown to facilitate GAS biofilm formation and virulence in a murine model of GAS respiratory infection (18). Recently, *in silico* analysis has revealed four AgI/II gene homologs in GBS, designated Group <u>B</u> *Streptococcus* <u>s</u>urface <u>p</u>roteins (BspA-D), that are distributed among GBS of different capsular serotypes and sequence types (15, 19, 20).

**Figure 1.**
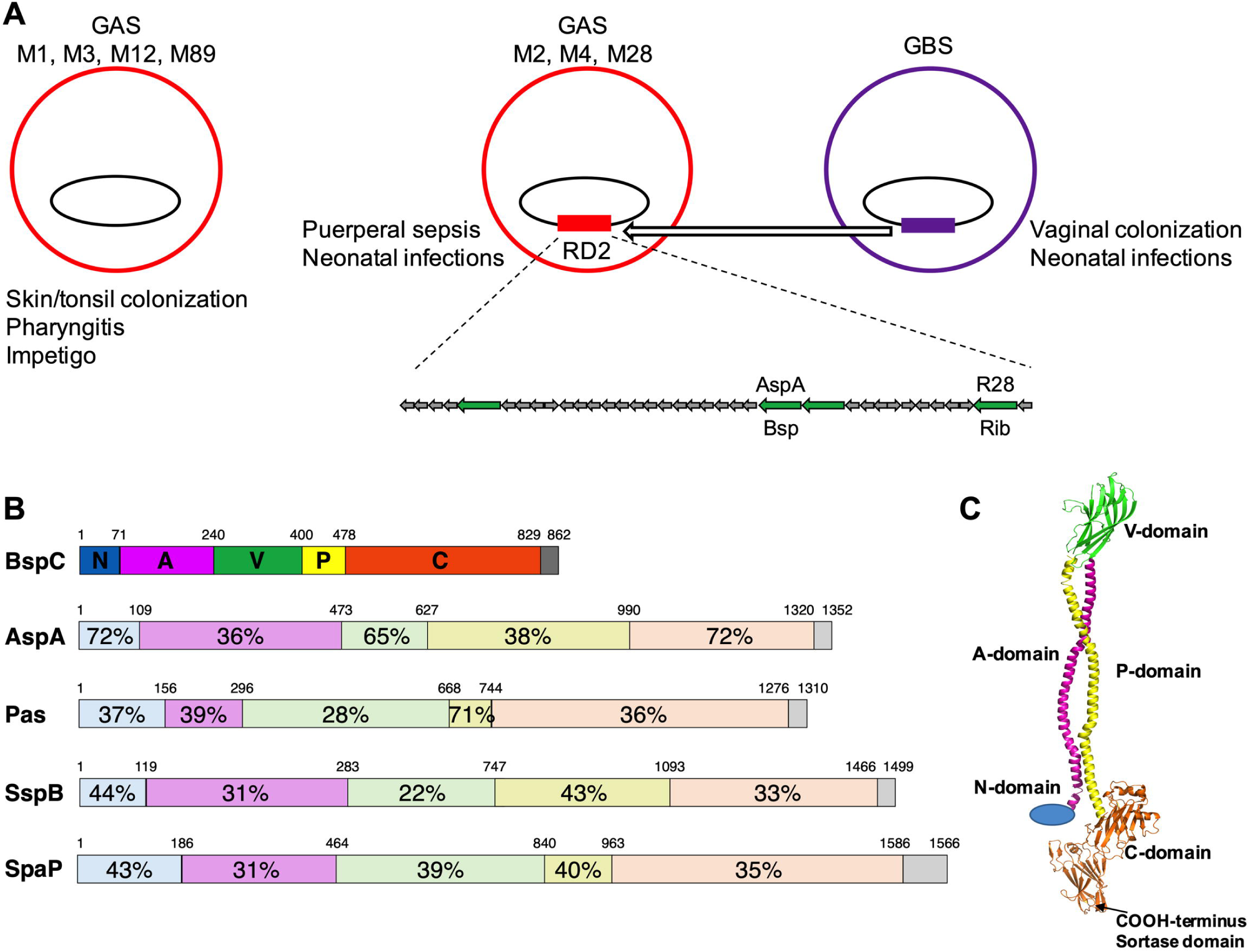
Analysis of BspC domain architecture (A) Diagram of MGAS6180 region of difference 2 (RD2). M2, M4, and M28 strains of Group A *Streptococcus* contain RD2, which was likely acquired from Group B *Streptococcus* through horizontal gene transfer and is absent in other common disease causing GAS *emm*-types. Open reading frames encoding LPXTG cell wall anchor domain-containing proteins are indicated with green arrows. Proteins AspA and R28 in GAS and their respective homologs in GBS, Bsp and Rib, have been previously described. The other two cell wall anchor domain-containing proteins have not been characterized. (B) Schematic of sequence homologies across antigen I/II family proteins from select *Streptococcus* species. Six structural regions are shown: N, N-terminal region; A, alanine-rich repeats; V, variable region; P, proline-rich repeats; C, C-terminal region; and the cell-wall anchor-containing region. Numbers indicate percentage amino acid residue identities to BspC from *S. agalactiae* strain COH1 (NCBI Ref. Seq.: WP_000277676.1). The proteins depicted are the antigen I/II proteins AspA from *S. pyogenes* strain M28 (WP_011285012.1), Pas from *S. intermedius* (WP_049476098.1), SspB from *S. gordonii* (WP_011999747), and SpaP from *S. mutans* (WP_024781655.1). (C) Hypothetical model of BspC generated using PyMOL.

Previous work has shown that BspA and BspB, which share 90% sequence identity, are found in GBS strain NEM316. BspA has been demonstrated to be important in biofilm formation as well as adherence to epithelial cells and may play a role in facilitating colonization through its ability to bind to vaginal epithelium as well as interact with the hyphal filaments of *Candida albicans* (15, 20), a frequent fungal colonizer of the lower female reproductive tract. Other GBS strains contain the homolog BspC, or in some cases BspD, which is over 99% identical to BspC, with the major difference being that BspD is missing the leader peptide for targeting to the cell surface by the Sec translocation machinery. While most of the variability between Bsp proteins is in the alanine-rich and proline-rich repeats, the V domain shares 96 to 100% identity across all Bsp homologs (15, 20). To date, there have not been any studies examining the impact of the AgI/II proteins on GBS invasive disease. A previous study by Chuzeville et al. identified 75 GBS genomes which contain an antigen I/II homolog. Of those 75 CDS, only 40 were associated with transcription and translation signals and out of those, 36 were 2952 base pairs in size encoding the full length BspC protein (19). Therefore, we chose to investigate the importance of the BspC antigen I/II homolog in the pathogenesis of meningitis. Using targeted mutagenesis, we show that BspC promotes adherence of bacteria to human cerebral microvascular endothelial cells (hCMEC) and interacts with the host cytoskeleton component, vimentin. Additionally, we found that BspC and vimentin contribute to the development of GBS meningitis in a mouse infection model. Lastly, we observe that BspC stimulates inflammatory signaling from brain endothelial cells *in vitro* and *in vivo* and that this immune signaling involves the NF-κB pathway.

## RESULTS

### Analysis of BspC domain architecture and construction of a *bspC* deletion strain

BspC contains all six domains characteristic of the AgI/II protein family, and shares high homology with other streptococcal AgI/II proteins, especially GAS AspA (Fig. 1B). The proposed domain organization of streptococcal AgI/II polypeptides comprises a stalk consisting of the α-helical A (alanine-rich repeats) domain and the polyproline II (PPII) helical P domain, separating the V (variable) domain and the C-terminal domain, which contains the LP*X*TG motif required for cell wall anchorage (14). While the GBS BspC structure is not known, the structure of several regions of the GBS homolog, BspA, has been solved (15). We generated a hypothetical model of full length BspC using PyMOL (The PyMOL Molecular Graphics System, Version 2.1 Schrödinger, LLC) for the purpose of showing the overall domain structure (Fig. 1C). Structures of individual BspC domains were generated using Phyre2 server (21). The V- and C-domains were modeled on the V- and C-domains of BspA (PDB entries 5DZ8 and 5DZA, respectively) (15), and approximately two-thirds of the A-domain sequence was modeled on human fibrinogen (PDB entry 3GHG) (22). It was not possible to generate models for the N- and P-domains, so the N-domain is shown as a sphere and the P-domain shown is a mirror image of the A-domain.

We performed precise in-frame allelic replacement to generate a Δ*bspC* mutant in GBS strain COH1, a hypervirulent GBS clinical isolate that is highly associated with meningitis (sequence type [ST]-17, serotype III) (23, 24), using a method as described previously (10). We further determined the presence of surface BspC expression in the WT and complemented strains compared to the Δ*bspC* mutant by flow cytometry and immunofluorescent staining with specific BspC antibodies (Fig. S1A-G). Growth curve analysis demonstrated that the Δ*bspC* mutant grew similarly to the WT parental strain under the conditions used here (Fig. S1H). Similarly, we observed no differences in hemolytic activity or capsule abundance between the WT and mutant strains (Fig. S1I-L). High-magnification scanning electron microscopy (SEM) images of COH1 strains showed that the Δ*bspC* deletion strain exhibits similar surface morphology to the isogenic wild type (Fig. 2A and 2B). However, lower-magnification SEM revealed that the Δ*bspC* mutant appeared to exhibit decreased interaction between neighboring cells (Fig. 2C,D).

**Figure 2.**
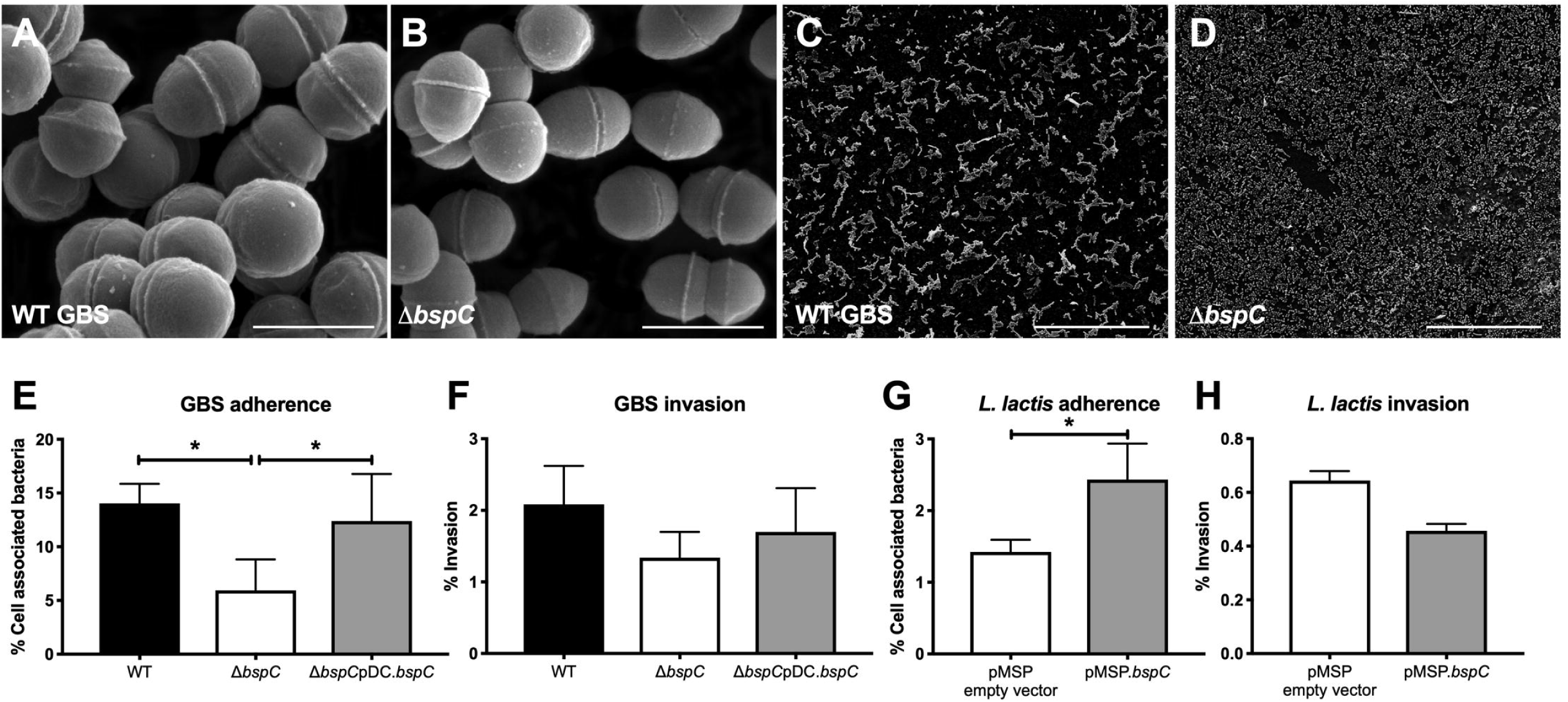
BspC bacterial adherence to endothelial cells *in vitro*. (A-D) Scanning electron microscopy images of WT (A and C) and Δ*bspC* mutant (B and D) GBS. Scale bar in high magnification images (A and B) is 1 μM. Scale bar in low magnification images (C and D) is 50 μM. (E) Adherence of WT GBS, the Δ*bspC* mutant, and the Δ*bspC*pDC.*bspC* complemented strain to hCMEC was assessed after a 30 min incubation. Total cell-associated bacteria are shown. (F) Invasion of WT GBS, the Δ*bspC* mutant, and the Δ*bspC*pDC.*bspC* strain was quantified after a 2 h infection. (G) Adherence and (H) invasion of *Lactococcus lactis* containing the pMSP empty vector or pMSP.*bspC* to hCMEC were quantified. Data indicates the percentage of the initial inoculum that was recovered. Experiments were performed three times with each condition in triplicate. Data from one representative experiment are shown and error bars represent the standard deviation. Statistical analysis: (E and F) One-way ANOVA with Tukey’s multiple comparisons test. (G and H) Unpaired t test. *, *P* < 0.05.

### BspC promotes bacterial adherence to endothelial cells *in vitro*

Since AgI/II proteins are known to demonstrate adhesive properties in other streptococci (14), we hypothesized that BspC would contribute to GBS interaction with brain endothelium. Thus, we characterized the ability of the Δ*bspC* mutant to attach to and invade hCMEC using our established adherence and invasion assays (10, 25). The Δ*bspC* mutant exhibited a significant decrease in adherence to hCMEC compared to WT GBS, and this defect was complemented when BspC was expressed in the Δ*bspC* mutant strain (Fig. 2E). This resulted in less recovery of intracellular Δ*bspC* mutant (Fig. 2F), but together these results indicate that BspC contributes primarily to bacterial attachment to hCMEC. To determine if BspC was sufficient to confer adhesion, we heterologously expressed the GBS *bspC* gene in the non-adherent, non-pathogenic bacterium *Lactococcus lactis*. Flow cytometric analysis of *L. lactis* confirmed surface expression of BspC protein in the strain containing the pMSP.*bspC* plasmid (Fig. S2). BspC expression resulted in a significant increase in *L. lactis* adherence to hCMEC compared to *L. lactis* containing the control vector, while invasion was not affected (Fig. 2G,H). These results demonstrate that BspC is both necessary and sufficient to confer bacterial adherence to hCMEC.

### BspC contributes to pathogenesis of GBS meningitis *in vivo*

Our results thus far suggest a primary role for BspC in GBS adherence to brain endothelium. We hypothesized that these *in vitro* phenotypes would translate into a diminished ability to penetrate the BBB and produce meningitis *in vivo*. Using our standard model of GBS hematogenous meningitis (10, 26, 27), mice were challenged with either WT GBS or the Δ*bspC* mutant as described in the Methods. The WT GBS strain caused significantly higher mortality than the isogenic Δ*bspC* mutant strain (Fig. 3A). By 48 hours, 80% of mice infected with WT COH1 had succumbed to death, while all of the mice infected with the Δ*bspC* mutant survived up to or past the experimental endpoint. In a subsequent experiment mice were infected with a lower dose of either WT GBS or the Δ*bspC* mutant and were sacrificed at a defined endpoint (48 hrs) to determine bacterial loads in blood, lung, and brain tissue. We recovered similar numbers of the Δ*bspC* mutant strain from mouse blood and lung compared to WT, however we observed a significant decrease in the amounts of the Δ*bspC* mutant recovered from the brain tissue (Fig. 3B). To confirm these results using other GBS strains, we constructed *bspC* gene deletions as described in the Methods in two other GBS strains: GBS 515 (ST-23, serotype Ia) and meningeal isolate 90356 (ST-17, serotype III). Mice infected with these mutant strains were also less susceptible to infection and exhibited decreased bacterial loads in the brain compared to the isogenic parental WT strain (Fig. S2). Interestingly, the Δ*bspC* mutant in the 515 GBS background, which is a different sequence type and serotype from the other two strains, appeared to also exhibit diminished infiltration into the mouse lungs.

**Figure 3.**
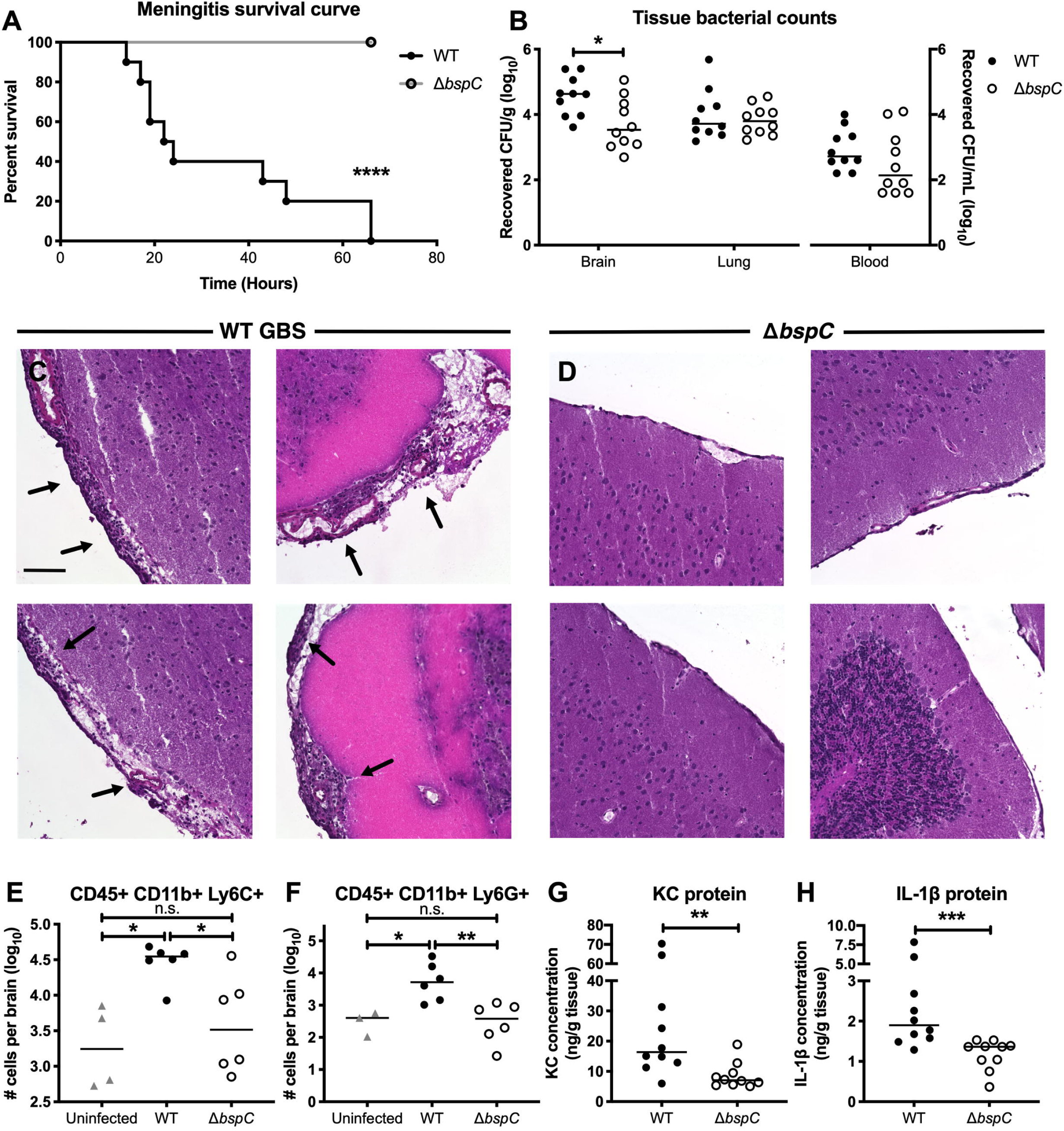
BspC contributes to pathogenesis of GBS meningitis *in vivo*. (A) A Kaplan-Meier plot showing survival of CD-1 mice infected with 10^9^ CFU of either WT GBS or the Δ*bspC* mutant. (B) 72 h after infection with 10^8^ CFU of either WT GBS or the Δ*bspC* mutant, mice were euthanized and bacterial loads in brain, lung, and blood were quantified. Lines indicate statistical median. (C and D) H&E images showing the leptomeninges on the surface of brains from CD-1 mice infected with either WT GBS (C) or the Δ*bspC* mutant (D). Arrows indicate areas of meningeal thickening and leukocyte infiltration. Scale bar is 100 μM. (E and F) Quantification of infiltrating immune cells in the brains of mice infected with WT GBS or the Δ*bspC* mutant. The presence CD45+, CD11b+, Ly6C+ (E), and CD45+, CD11b+, Ly6G+ (F) cells was determined using flow cytometry. (G and H) ELISA was performed on mouse brain tissue homogenates to assess cytokine protein levels. KC (G) and IL-1β (H) were quantified for brains from mice challenged with either WT GBS or the Δ*bspC* mutant. Statistical analysis: (A) Log-rank test. (B) Two-way ANOVA with Sidak’s multiple comparisons test. (E and F) One-way ANOVA with Sidak’s multiple comparisons test. (G and H) Mann-Whitney test. *, *P* < 0.0005; **, *P* < 0.005; ***, *P* < 0.0005; ****, *P* < 0.00005.

As excessive inflammation is associated with CNS injury during meningitis, we performed histological analysis of brains from infected animals. In WT infected mice, we observed leukocyte infiltration and meningeal thickening characteristic of meningitis that was absent in the mice infected with the Δ*bspC* mutant strain (Fig. 3C,D). Representative images from 8 mice, 4 infected with WT (Fig. 3C) and 4 infected with the Δ*bspC* mutant (Fig. 3D) are shown where the major areas of inflammation were observed. In subsequent experiments to quantify the total inflammatory infiltrate, whole brains from uninfected mice and mice infected with either WT GBS or Δ*bspC* mutant GBS were processed and analyzed by flow cytometry as described in the Methods. There was no significant difference in the numbers of CD45 positive cells between the groups of mice, however within the CD11b positive population we observed higher numbers of Ly6C positive and Ly6G positive cells in the brains of animals infected with WT GBS compared to uninfected mice and mice infected with the Δ*bspC* mutant strain (Fig. 3E,F), indicating an increased population of monocytes and neutrophils. Consistent with these results, we further observed that mice challenged with WT GBS had significantly more of the neutrophil chemokine, KC, as well as the proinflammatory cytokine, IL-1β, in brain homogenates than Δ*bspC* mutant infected animals. (Fig 3G, H)

### BspC is necessary and sufficient to induce neutrophil chemokine signaling

To further characterize the role of BspC in stimulating immune signaling pathways we infected hCMEC with WT GBS, the Δ*bspC* mutant, or the complemented strain. After four hours of infection, we collected cells and isolated RNA for RT-qPCR analysis to quantify IL-8 and CXCL-1 transcripts. We focused on the neutrophil chemokines IL-8 and CXCL-1 as these cytokines are highly induced during bacterial meningitis (28) and we observed an increase in neutrophilic infiltrate in brain tissue during the development of GBS meningitis in our mouse model. Cells infected with WT GBS had significantly increases of both transcripts compared to cells infected with the Δ*bspC* mutant. Complementation of the *bspC* mutation restored the ability of the bacteria to stimulate the expression of IL-8 and CXCL-1 (Fig. 4A,B). Additionally, treatment of hCMEC with purified BspC protein resulted in increased transcript abundance (Fig. 4,D) and protein secretion (Fig. 4E,F) for both IL-8 and CXCL-1 compared to untreated cells or treatment with control protein, CshA from *Streptococcus gordonii,* that was similarly purified from *E. coli*.

**Figure 4.**
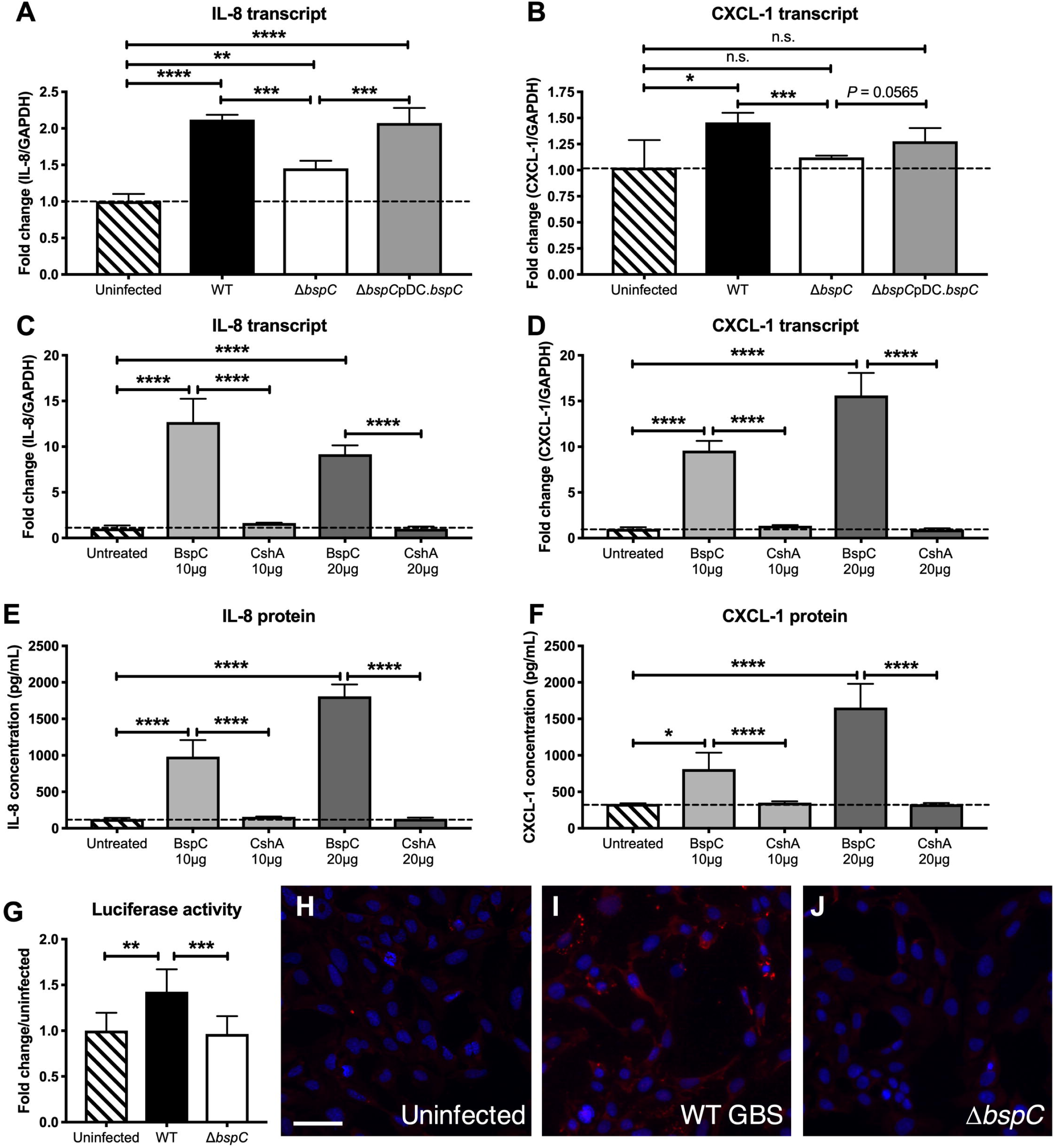
BspC is necessary and sufficient to induce neutrophil chemokine signaling. (A and B) RT-qPCR to assess IL-8 (A) and CXCL-1 (B) transcript levels in uninfected hCMEC or hCMEC infected with WT GBS, the Δ*bspC* mutant, or the complemented strain. (C and D) RT-qPCR analysis quantifying IL-8 (C) and CXCL-1 (D) transcripts in hCMEC treated with purified BspC protein or the *S. gordonii* CshA protein. (E and F) ELISA to measure IL-8 (E) and CXCL-1 (F) protein secretion by hCMEC treated with BspC or CshA. (G) Luciferase activity of uninfected Hela-57A cells and cells infected with WT GBS or the Δ*bspC* mutant was assessed. Experiments were performed at least three times with each condition in triplicate. Data from representative experiments are shown and error bars represent the standard deviation of the mean for one experiment. Data were analyzed with One-way ANOVA with Sidak’s multiple comparisons test. (H-J) Immunofluorescent staining showing p65 localization in uninfected hCMEC (H), hCMEC infected with WT GBS (I), and hCMEC infected with the Δ*bspC* mutant (J). Following infection, cells were fixed with formaldehyde and incubated with a rabbit antibody to p65. Nuclei were labelled with DAPI. Scale bar is 50 μM. *, *P* < 0.05; **, *P* < 0.005; ***, *P* < 0.0005; ****, *P* < 0.00005.

Nuclear factor-κB (NF-κB) represents a family of inducible transcription factors, which regulates a large array of genes involved in different processes of the immune and inflammatory responses, including IL-8, CXCL-1 and IL-1 (29). To assess whether the NF-κB pathway is activated by BspC, we utilized the Hela-57A NF-κB luciferase reporter cell line as described previously (30). Cells infected with WT GBS had significantly higher luciferase activity than uninfected and Δ*bspC* mutant GBS infected cells, indicating that BspC contributes NF-κB activation (Fig. 4G). Additionally, immunofluorescent staining of hCMEC revealed an increase in P65 expression, an indicator of NF-κB activation, during infection with WT GBS but not in response to infection with Δ*bspC* mutant GBS (Fig. 4H-J).

### BspC interacts with the host endothelial cytoskeletal component vimentin

We next sought to identify the host protein receptor on brain endothelial cells that interacts with BspC. Membrane proteins of hCMEC were separated by 2-dimensional electrophoresis (2-DE), then blotted to a PVDF membrane. Following incubation with biotinylated BspC protein, the PVDF membrane was incubated with a streptavidin antibody conjugated to HRP. While many proteins were detected on the Coomassie stained gel, biotinylated BspC protein specifically interacted predominately with one spot in the PVDF membrane with molecular mass around 50-55 kDa and isoelectric point (pI) around 5 (Fig. 5A,B). The corresponding spot from the Coomassie-stained 2-DE gel was excised and digested with trypsin (Fig. 5C). Resulting peptides were analyzed by liquid chromatography-tandem mass spectrometry. The spectra from the spot yielded 158 peptide sequences which matched to human vimentin. The molecular weight, 53.6 kDa, and calculated pI, 5.12, of vimentin match the values for the spot on the 2-DE gels. This procedure was repeated for membrane proteins from another human brain endothelial cell line (hBMEC) that has been used previously to study GBS interactions (31). Mass spectrometry analysis also determined that BspC interacted with human vimentin (Fig. S3A-C). The control far Western blot with the streptavidin antibody conjugated to HRP did not show any hybridization (Fig. S3D).

**Figure 5.**
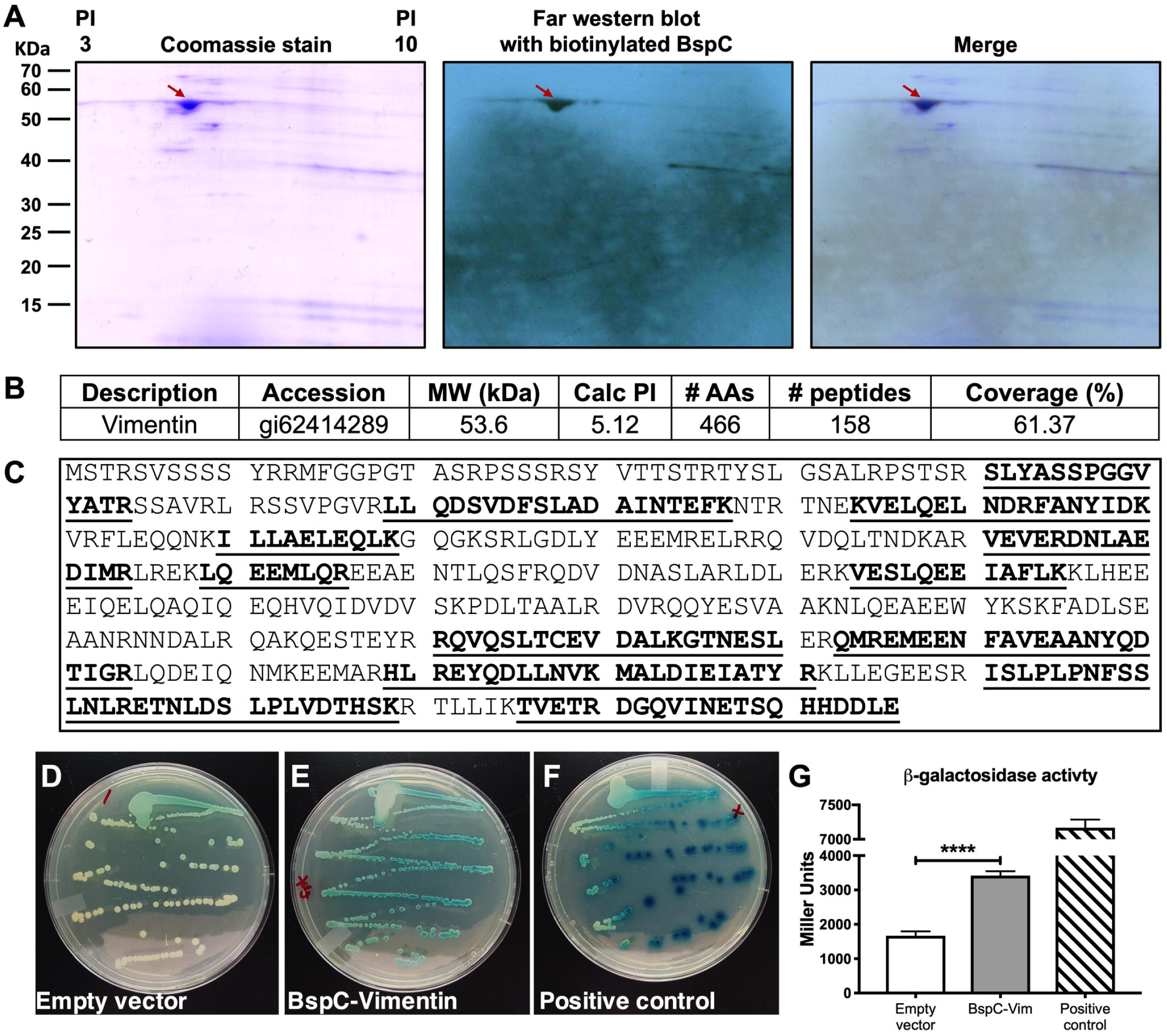
BspC interacts with the host endothelial cytoskeletal component vimentin. (A) Far Western blot analysis of hCMEC membrane proteins probed with biotinylated BspC. Membrane proteins were extracted from hCMEC and separated by 2-DE in duplicate. One gel was transferred to a PVDF membrane and probed with biotinylated BspC. The specific interaction of BspC was detected by a streptavidin antibody conjugated to HRP and visualized by x-ray film exposure. The other gel was stained with Coomassie blue. The spot identified from the x-ray film was aligned to the Coomassie stained gel and the corresponding spot was excised and digested with trypsin. (B) Electrospray ionization-tandem mass spectrometry analysis identifies the protein vimentin. (C) Amino acid sequence of human vimentin. Underscored and bolded are the peptide sequences identified from MS analysis. (D-F) Bacterial two-hybrid assay using *E. coli* containing empty vector controls (D), the BspC and vimentin vectors (E), and the positive control vectors (F). (G) β-galactosidase activity of *E. coli* was quantified using a Miller assay. The Miller assay was performed three times with each condition in five replicates. Data from one representative experiment is shown, error bars represent the standard deviation of the mean. Data were analyzed using one-way ANOVA with Dunnett’s multiple comparisons test. ****, *P* < 0.00005.

To confirm these protein-protein interactions *in vivo*, we employed a bacterial two-hybrid system (BACTH, “Bacterial Adenylate Cyclase-Based Two-Hybrid”) (32). This system is based on the interaction-mediated reconstitution of a cyclic adenosine monophosphate (cAMP) signaling cascade in *Escherichia coli*, and has been used successfully to detect and analyze the interactions between a number of different proteins from both prokaryotes and eukaryotes (33). Using a commercially available kit (Euromedex) according to manufacturer’s directions and as described previously (34), vimentin was cloned and fused to the T25 fragment as a N-terminal fusion (T25-Vimentin), using the pKT25 plasmid, and *bspC* was cloned as a c-terminal fusion (BspC-T18) using the pUT18 plasmid. To test for interaction these plasmids were transformed into an *E. coli* strain lacking adenylate cyclase (*cyaA*). We observed blue colonies when grown on LB agar plates containing X-gal, indicating β-galactosidase activity and a positive interaction between Vimentin and BspC compared to the empty vector control (Fig. 5D,E). A leucine zipper that is fused to the T25 and T18 fragments served as the positive control for the system (Fig. 5F). To quantify β-galactosidase activity, cells grown to log phase were permeabilized with 0.1 % SDS and toluene and enzymatic activity measured by adding ONPG as described previously (34). The *E. coli* strain containing both vimentin and BspC expressing plasmids exhibited increased β-galactosidase activity (Miller units) compared to the empty vector control strain (Fig. 5G). We also quantified the interaction between BspC and vimentin by performing microscale thermophoresis (MST) (35) as described in Methods. The dissociation constant (*K*_d_) was 3.39μM as calculated from the fitted curve that plots normalized fluorescence against concentration of vimentin (Fig. S4E). These results demonstrate a direct interaction between BspC and vimentin.

### GBS co-localization with cell-surface vimentin is dependent on BspC

To visualize the localization of WT GBS and vimentin in brain endothelial cells we performed imaging flow cytometry of uninfected and WT GBS infected hCMEC. Cells were fixed, permeabilized, and incubated with antibodies to vimentin and GBS. We observed that vimentin protein was present throughout the cytoplasm of uninfected hCMEC (Fig. 6A,B), but during infection GBS co-localized with vimentin near the surface of infected cells (Fig. 6C,D). To further characterize the localization of GBS and vimentin, we performed immunofluorescent staining of hCMEC infected with either WT GBS or the Δ*bspC* mutant. Following infection, cells were fixed but not permeabilized to permit labeling of only extracellular bacteria and surface expressed vimentin. We observed that surface vimentin of hCMEC co-localized with WT GBS bacteria, while this was not seen for hCMEC infected with the Δ*bspC* mutant (Fig. 6E-H). To quantify co-localization of GBS with vimentin, the number of bacteria that overlapped with the vimentin signal was divided by the total number of bacteria in each field of view (Fig. 6I). No staining of either GBS or vimentin was observed in IgG controls (Fig 6J-L). Additionally, as a control, we did not observe staining of GBS with the anti-vimentin antibody (data not shown).

**Figure 6.**
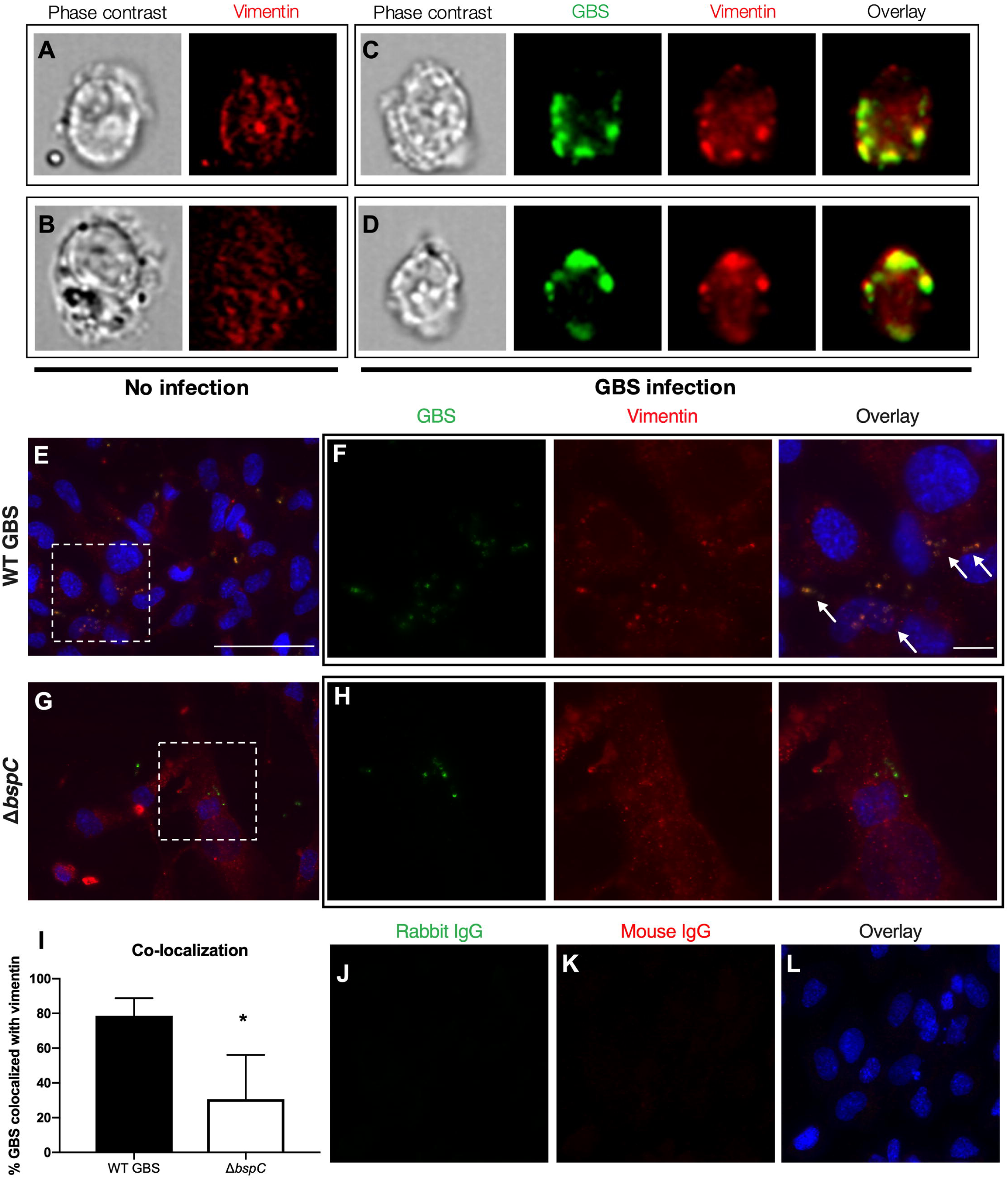
GBS co-localization with cell-surface vimentin is dependent on BspC. (A-D) Imaging flow cytometry showing vimentin localization in uninfected hCMEC (A and B) and hCMEC infected with WT GBS (C-D). Cells were fixed and permeabilized prior to staining with antibodies to vimentin and the group B carbohydrate antigen to label cell-associated GBS. (E-H) Immunofluorescent staining of hCMEC monolayers infected with WT GBS (E-F) or the Δ*bspC* mutant (G-H). Following a one-hour infection, hCMEC were washed to remove nonadherent bacteria then fixed and labelled with antibodies to vimentin and GBS. Nuclei were labelled with DAPI. Magnified images of the areas highlighted in (E) and (G) are shown in (F) and (H). Scale bar in (E) is 50 μm. Scale bare in (F) is 10 μm. Arrows indicate GBS co-localizing with vimentin. (I) Quantification of co-localization of GBS and vimentin was performed by dividing the number of GBS that co-localize with vimentin by the total number of GBS. Error bars represent the median. Statistical analysis was performed using and unpaired t test. *, *P* < 0.05. (J-L) Negative staining controls of hCMEC monolayers infected with WT GBS.

### BspC and vimentin promote GBS attachment to cells *in vitro*

To first determine if BspC-mediated attachment to cells is dependent on vimentin, we infected HEK293T cells with lentiviruses containing either the vimentin expression plasmid pLenti-VIM or the vector control pLenti-mock. Immunofluorescent staining reveals that HEK293T pLenti-mock cells do not express vimentin while the HEK203T pLenti-VIM clone exhibits strong vimentin labelling (Fig. 7A,B). WT GBS was significantly more adherent to HEK293T cells that express vimentin while the Δ*bspC* mutant showed no difference in attachment to either cell line (Fig. 7C). Next, we assessed the effect of blocking the vimentin-GBS interaction by treating hCMEC with anti-vimentin antibodies prior to infection with GBS (Fig. 7D). Treatment with a vimentin antibody that recognizes the N-terminal epitope (AA31-80), as well as with IgG isotype controls did not alter adherence of the WT or Δ*bspC* mutant strains, however pre-incubation with the mouse V9 antibody (36, 37), which reacts with the C-terminal of vimentin (AA405-466), reduced WT GBS adherence to levels comparable to the adherence of the Δ*bspC* mutant (Fig. 7E,F). These results indicate that the interaction between BspC and cell-surface vimentin is dependent on the C-terminus of vimentin.

**Figure 7.**
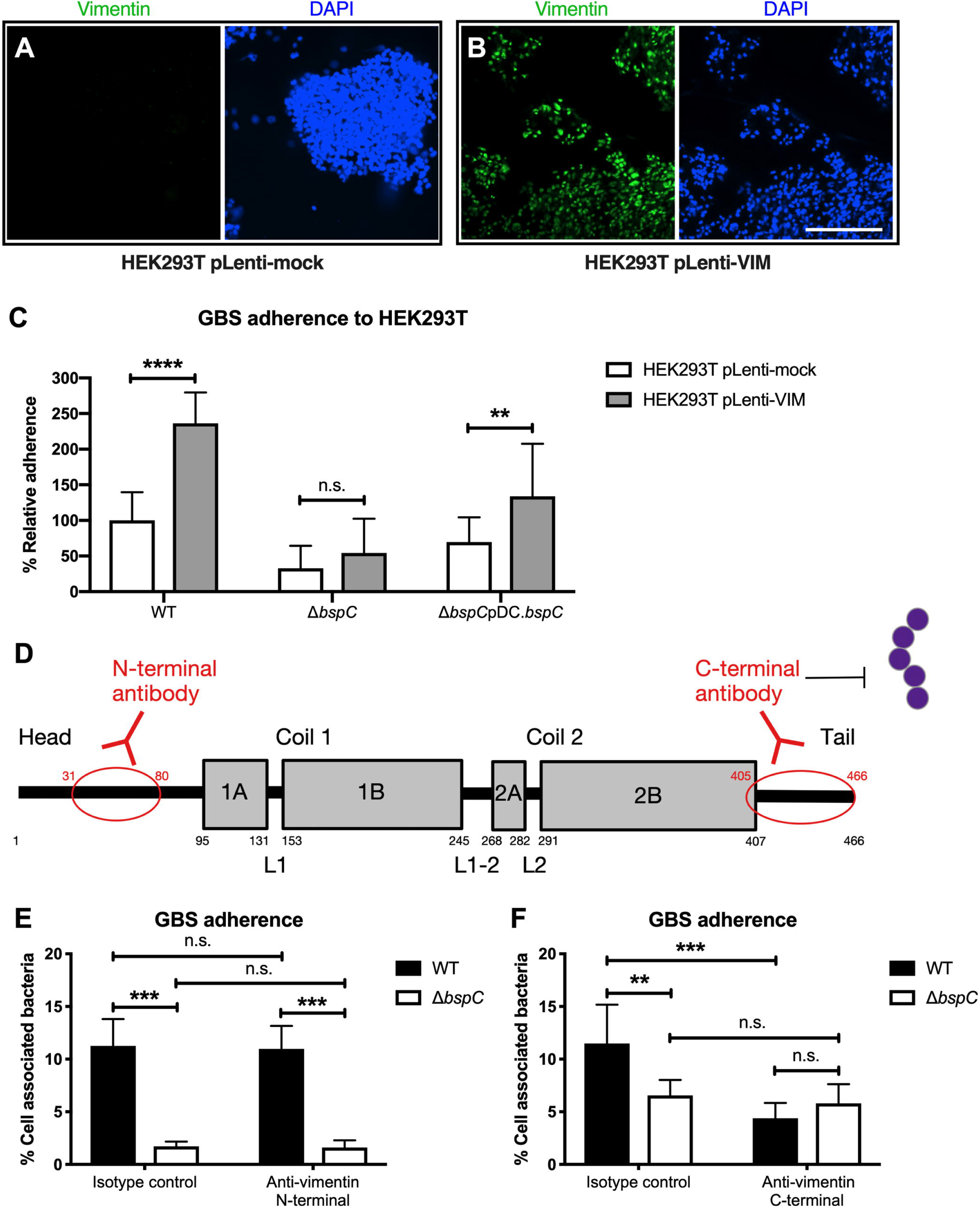
GBS adherence to cells is dependent on vimentin. (A and B) Immunofluorescent staining to show vimentin expression of HEK293T pLenti-Mock (A) and HEK293T pLenti-VIM (B) cells. Scale bar is 200 μm. (C) Adherence of WT GBS, the Δ*bspC* mutant, and the complemented strain to transfected HEK293T cells. Data from two independent experiments with each condition in 8 replicates is combined. (D) Schematic showing regions of vimentin recognized by the monoclonal antibodies. (E and F) hCMEC were pre-incubated with antibodies to the N-terminal (E) or the C-terminal (F) of vimentin or with an IgG control prior to a 30 min infection with either WT GBS or the Δ*bspC* mutant. Experiments were performed three times with each condition in triplicate and data from one representative experiment are shown. Error bars represent the standard deviation of the mean. Statistical analysis was performed using a two-way ANOVA with Sidak’s multiple comparisons test. **, *P* < 0.005; ***, *P* < 0.0005; ****, *P* < 0.00005.

### Vimentin contributes to the pathogenesis of meningitis *in vivo*

We obtained WT 129 and 129 vimentin KO mice and confirmed the absence of vimentin in the brain endothelium of the KO animals by immunofluorescent staining (Fig. S5A,B). To determine the necessity of vimentin in GBS meningitis disease progression, we infected WT and vimentin KO mice with WT GBS and observed that they were less susceptible to GBS infection and exhibited increased survival compared to WT animals (Fig. 8A). Further, significantly less bacteria were recovered from the tissues of KO mice compared to the tissues of WT mice (Fig. 8B). WT and vimentin KO mice infected with Δ*bspC* mutant GBS showed no difference in survival and tissue bacterial counts (Fig. S6). Additionally, for animals infected with WT GBS, we detected significantly less KC and IL-1β in brain tissues of vimentin KO compared to WT mice, suggesting that vimentin contributes to the initiation of immune signaling pathways during GBS infection (Fig. 8C,D). Taken together these results indicate the importance of vimentin in GBS dissemination into the brain and meningitis disease progression.

**Figure 8.**
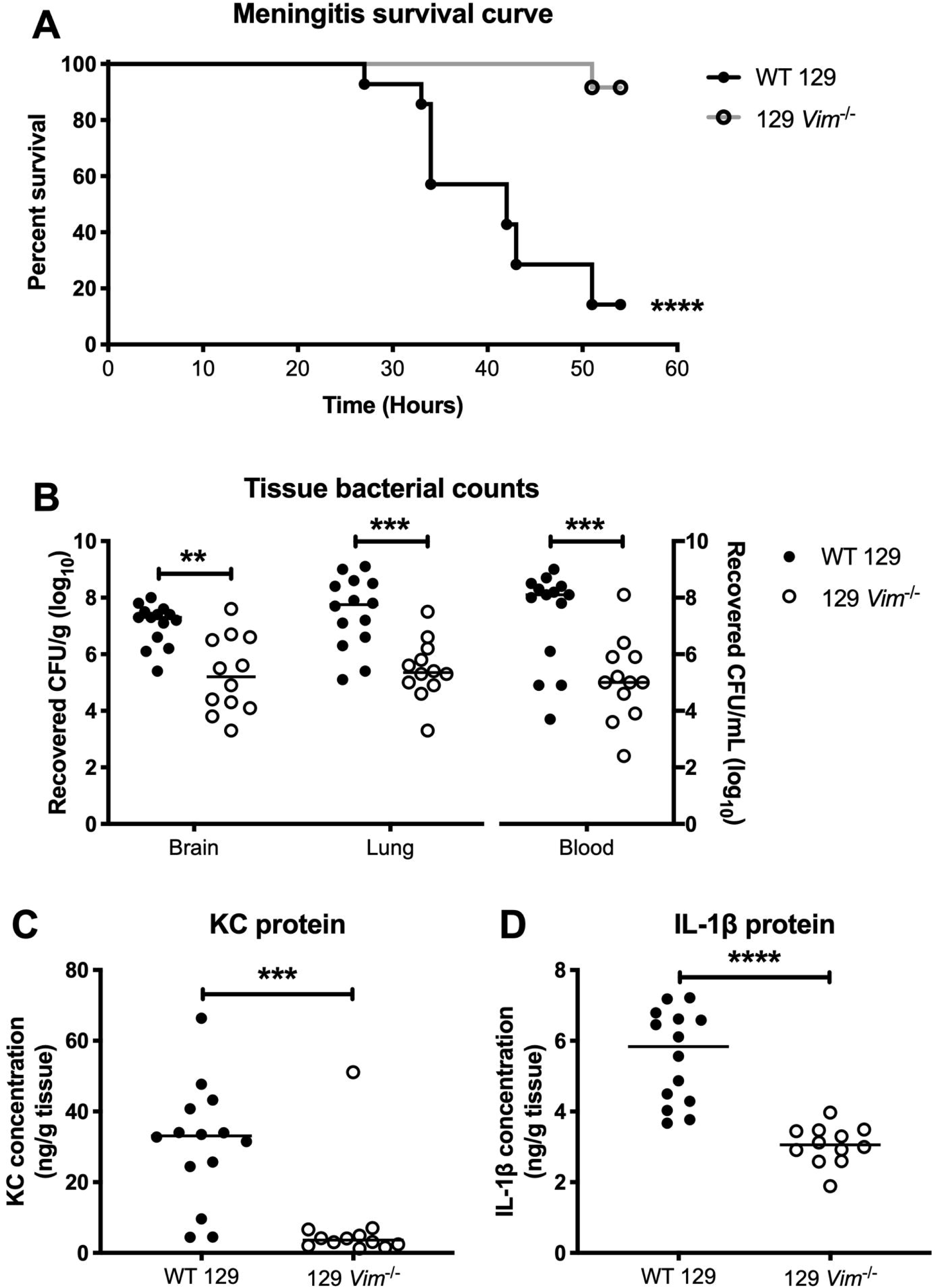
Vimentin contributes to the pathogenesis of GBS infection. (A) Kaplan-Meier plot showing survival of WT 129 and 129 *Vim*^-/-^ mice infected with 10^8^ CFU of WT GBS. (B) Tissue bacterial counts for WT 129 and 129 *Vim*^-/-^ mice infected with WT GBS. (C and D) ELISA to quantify KC (C) and IL-1β (D) protein in brain tissue homogenates. Statistical analysis: (A) Log-rank test. (B-D) Mann-Whitney test. **, *P* < 0.005; ***, *P* < 0.0005; ****, *P* < 0.00005.

**Figure 9.**
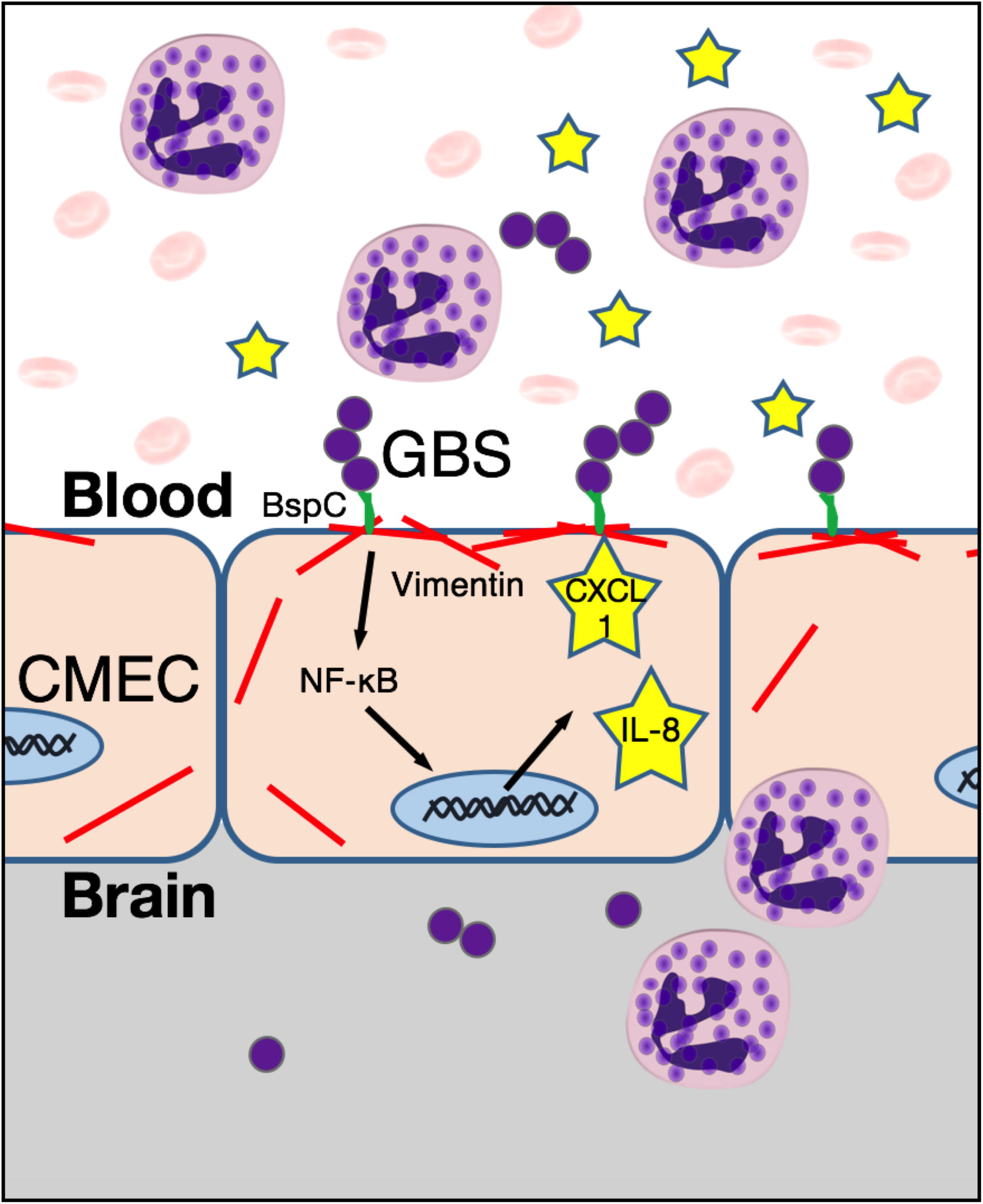
Summary of the role of the BspC-vimentin interaction in promoting meningitis. BspC interacts with vimentin on the surface of CMEC to promote GBS attachment and the production of the neutrophil recruiting cytokines IL-8 and CXCL-1 through the NF-κB pathway.

## DISCUSSION

Our studies reveal a unique requirement for the Group B streptococcal Antigen I/II protein, BspC, to brain penetration by GBS, the leading agent of neonatal bacterial meningitis. A decreased ability by the GBS Δ*bspC* mutant to attach to brain endothelium and induce neutrophil chemoattractants *in vitro* was correlated with a reduced risk for development of meningitis and markedly diminished lethality *in vivo*. We identified that BspC interacts directly with host vimentin and that blocking this interaction abrogated BspC-mediated attachment to hCMEC. Further, vimentin deficient mice infected with GBS exhibited decreased mortality, bacterial brain loads, and cytokine production in brain tissue. These results corroborate the growing evidence that this intermediate filament protein plays important roles in the pathogenesis of bacterial infections (38), and provide new evidence for the pivotal role of the BspC adhesin in GBS CNS disease (Fig. 8).

The oral streptococcal AgI/II adhesins range in composition from 1310 - 1653 amino acid (AA) residues, while GBS AgI/II proteins are smaller (826-932 AA residues) (39). The primary sequences of AgI/II proteins are comprised of six distinct regions (Fig. 1B), several of which have been shown to mediate the interaction to various host substrates. The *S. mutans* protein SpaP as well as the *S. gordonii* proteins SspA and SspB have been demonstrated to interact specifically with the innate immunity scavenger protein gp-340. (40) Recently, the GAS AgI/II protein AspA, as well as BspA, the AgI/II homolog expressed mainly by the GBS strain NEM316, have also been shown to bind to immobilized gp-340 (15, 18). Gp-340 proteins are involved in various host innate defenses and are present in mucosal secretions, including saliva in the oral cavity and bronchial alveolar fluid in the lung. They can form complexes with other mucosal components such as mucins and function to trap microbes for clearance. However, when gp-340 is immobilized, it can be used by bacteria as a receptor for adherence to the host surface. (15, 41-44). There is evidence that the Variable (V) regions of SspB, SpaP, AspA and BspA facilitate gp340-binding activity (15, 45-47). AgI/II family adhesins have also been shown to interact with other host factors including fibronectin, collagen, and β1 integrins to promote host colonization (14, 48-50), demonstrating the multifactorial nature of these adhesins. It is unknown if GBS BspC interactions with other host factors are similar to those of the AgI/II proteins from other streptococci, particularly since the respective V-domains of these homologs are distant enough to suggest different binding partners. Previous work by Chuzeville *et al*. suggests that the integrative and conjugative element which contains the *bspC* gene can contribute to bacterial adherence to fibrinogen (19). Our MST experiments reveal the *K*_d_ for the interaction between BspC and vimentin to be 3.39 μM. This binding affinity is very similar to that observed for other multifunctional bacterial adhesins and their various host ligands. For example, the *K*_d_ for the interaction between fibronectin-binding protein B of *Staphylococcus aureus* and fibrinogen, elastin, and fibronectin has been demonstrated to be 2 μM, 3.2 μM, and 2.5 μM, respectively (51). Whether BspC can promote adherence to other host factors requires further investigation as these interactions may be critical to GBS colonization of mucosal surfaces such as the gut and the vaginal tract.

Here we show that GBS BspC interacts with host vimentin, an important cytoskeletal protein belonging to class III intermediate filaments. Vimentin is located in the cytoplasm and functions as an intracellular scaffolding protein that maintains structural and mechanical cell integrity (52). However, vimentin is also found on the surface of numerous cells such as T cells, platelets, neutrophils, activated macrophages, vascular endothelial cells, skeletal muscle cells and brain microvascular endothelial cells (53–60). Vimentin also mediates a variety of cell processes including cell adhesion, immune signaling, and autophagy (55, 61, 62). Further, the role of cell surface vimentin as an attachment receptor facilitating bacterial or viral entry, has been previously documented for other pathogens (38, 63-65). The BspC domain that mediates the vimentin interaction is currently under investigation. As the V domain is likely projected from the cell surface and has been implicated in host interactions for other streptococci, we hypothesize that this may be a critical domain for this interaction. Additionally, as the V-domain of other Bsp homologs share 96-100% identity with the V-domain of BspC, we predict that the other Bsp proteins might also interact with vimentin, but this would be a topic for future investigation.

There is a growing body of evidence that various bacteria can interact with vimentin to promote their pathogenesis, including *Escherichia coli* K1, *Salmonella enterica*, *Streptococcus pyogenes*, and *Listeria monocytogenes.* Thus, vimentin has been shown to be important in experimental models of infection at body sites other than the brain (38, 57, 60, 66, 67). Interestingly, previous studies on the meningeal pathogen *E. coli* K1 have demonstrated that the bacterial surface factor, IbeA, interacts with vimentin to promote bacterial uptake into brain endothelial cells (60, 68). Similarly, while our study was underway, it was reported that another bacterium capable of causing meningitis, *L. monocytogenes,* uses InlF to interact with vimentin to promote brain invasion (67). Along with our results presented here, this may suggest a common mechanism for meningeal bacterial pathogens to penetrate the BBB and cause CNS disease. However, our analysis of these three bacteria proteins showed no homology or predicted regions that might commonly interact with vimentin. Furthermore, the interaction between the *E. coli* receptor IbeA and the *L. monocytogenes* receptor InlF with cell-surface vimentin can be blocked by an antibody to the N-terminal region of vimentin (60, 67), while we demonstrate that the interaction between BspC and cell-surface vimentin can be blocked with an antibody to the C-terminal of vimentin. The implications of this unique interaction between a bacterial receptor and the C-terminal of vimentin remain to be explored.

Neuronal injury during bacterial meningitis involves both microbial and host factors, and subsequent to attachment to the brain endothelium and penetration of the BBB, GBS stimulation of host immune pathways is the next important step in the progression of meningitis. The release of inflammatory factors by brain endothelial cells, microglia, astrocytes, and infiltrating immune cells can exacerbate neuronal injury (9). Our data suggest that, like other streptococcal AgI/II family polypeptides, BspC plays a role in immune stimulation. AgI/II family proteins contain two antigenic regions (the antigens I/II and II) (69) and this ability to elicit an inflammatory response makes SpaP, the *S. mutans* AgI/II protein, an attractive candidate for vaccine development to prevent dental caries (14, 70). In this study we found that BspC can stimulate NF-κB activation and the release of the proinflammatory cytokines IL-8 and CXCL-1 from hCMEC. Both of these chemokines are major neutrophil recruiting chemoattractants and are the most highly induced during GBS infection (27, 71). We observed that mice infected with GBS mutants that lack BspC exhibited lower brain bacterial loads and less meningeal inflammation compared to animals challenged with WT GBS. Interestingly, WT and Δ*bspC* mutant bacterial loads were similar in the blood, indicating that BspC may not influence GBS survival and proliferation in the blood; however further investigation is warranted.

This study demonstrates for the first time the importance of a streptococcal AgI/II protein, BspC, in the progression of bacterial meningitis. Our data demonstrate that BspC, likely in concert with other GBS surface determinants mentioned above (pili, Srr1/2, SfbA), contributes to the critical first step of GBS attachment to brain endothelium. As the other described GBS surface factors have been shown to interact with ECM components, BspC may mediate a more direct interaction with the host cell as it facilitates interaction with vimentin. We have observed a unique requirement for vimentin to the pathogenesis of CNS disease; vimentin KO mice were markedly less susceptible to GBS infection and exhibited reduced bacterial tissue load and inflammatory signaling. Vimentin is also known to act as a scaffold for important signaling molecules and mediates the activation of a variety of signaling pathways including NOD2 (nucleotide-binding oligomerization domain-containing protein 2) and NLRP3 (nucleotide-binding domain, leucine-rich-containing family pyrin domain-containing-3) that recognize bacterial peptidoglycan and activate inflammatory response via NF-κB signaling (68, 72, 73). Thus, continued investigation into the mechanisms of how BspC-vimentin interactions dually promote bacterial attachment and immune responses, as well as how BspC expression may be regulated and whether known GBS two-component systems are involved, is warranted. These studies will provide important information that may inform future therapeutic strategies to limit GBS disease progression.

## MATERIALS AND METHODS

### Ethics statement

Animal experiments were approved by the committee on the use and care of animals at San Diego State University (SDSU) protocol #16-10-021D and at University of Colorado School of Medicine protocol #00316 and performed using accepted veterinary standards. San Diego State University and the University of Colorado School of Medicine are AAALAC accredited; and the facilities meet and adhere to the standards in the “Guide for the Care and Use of Laboratory Animals”.

### Bacterial strains, growth conditions, proteins, and antibodies

GBS clinical isolate COH1 (serotype III) (74), 515 (serotype Ia) (20), the recent meningitis isolate 90356 (serotype III) (75) and their isogenic Δ*bspC* mutants were used for the experiments. GBS strains were grown in THB (Hardy Diagnostics) at 37°C, and growth was monitored by measuring the optical density at 600 nm (OD_600_). *Lactococcus lactis* strains were grown in M17 medium (BD Biosciences) supplemented with 0.5% glucose at 30°C. For antibiotic selection, 2 μg/mL chloramphenicol (Sigma) and 5 μg/mL erythromycin (Sigma) were incorporated into the growth medium. BspC and CshA recombinant proteins, and the BspC antibody were purified as described previously (15, 76). The anti-BspC polyclonal antibody was further adsorbed (as described in (77)) against COH1Δ*bspC* bacteria to remove natural rabbit antibodies that react with bacterial surface antigens. Briefly, anti-BspC antibody was diluted to 2.28 mg/mL in PBS and incubated with COH1Δ*bspC* bacteria overnight at 4°C, with rotation. Bacteria were pelleted by centrifugation and the supernatant was collected and filtered using 0.22 µM cellulose acetate SpinX centrifuge tube filters (Costar). A normal rabbit IgG antibody (Invitrogen) was adsorbed as described above, and utilized as a negative isotype control.

### Targeted mutagenesis and *complementation* vector construction

The Δ*bspC* mutant was generated in COH1and 90356 by in-frame allelic replacement with a chloramphenicol resistance cassette by homologous recombination using a method previously described (10). A knockout construct was generated by amplifying up- and down-flanking regions of the *bspC* gene from COH1 genomic DNA using primer pairs of 5’flank-F (GCAGACACCGATTGCACAAGC)/R (GAAGGCGATCTTGCCCTCAA) and 3’flank-F (GTCAGCTATCGGTTTAGCAGG)/R(CTATACACGCCTACAGGTGTC). The chloramphenicol resistance (*cat*) cassette was amplified with primers Cat-F (GAGGGCAAGATCGCCTTCATGGAGAAAAAAATCACTGGAT) and Cat-R (CTGCTAAACCGATAGCTGACTTACGCCCCGCCCTGCCACT). Then the construct of two flanks along with the *cat* cassette was amplified with a pair of nest primers, Nest-xhoI (CCFCTCGAGGATGCTCAAGATGCACTCAC) and Nest-xbaI (GCTCTAGACGAGCCAAATTACCCCTCCT), which was then cloned into the pHY304 vector (78) and propagated in *E. coli* strain DH5α (79) prior to isolation and transformation to COH1 and 90356 GBS. A Δ*bspC* mutant had been generated previously in 515 (20). The complemented strain of Δ*bspC* mutant in COH1 was generated by cloning *bspC* into pDCerm, an *E. coli*-GBS shuttle expression vector. Gene *bspC* was amplified from GBS 515 genomic DNA using primers pDC.*bspC*.F (TGGGTACCAGGAGAAAATATGTATAAAAATCAAAC) and pDC.*bspC*.R (CCGGGAGCTCGCAGGTCCAGCTTCAAATC), designed to encode a *Kpn*I and *Sac*I restriction site at its termini respectively. This *bspC* amplicon was then cloned into pDCerm and propagated in *E. coli* strain Stellar^TM^ (ClonTech), prior to isolation and transformation into COH1 GBS. A *L. lactis* strain expressing BspC had been generated previously (20).

### Hemolysis assay

GBS strains were grown to an OD_600_ of 0.4, harvested by centrifugation, and resuspended in PBS. A total of 1 x 10^8^ CFU was added to fresh sheep blood (VWR) in V-bottom 96-well plates (Corning). The plates were sealed and incubated at 37°C with agitation for 1 h. The plates were centrifuged at 200 x *g* for 10 min, and 100 μl of the supernatant was transferred to a flat-bottom 96-well plate. The absorbance at 541 (*A*_541_) was read, and percent hemolysis was calculated by comparing the *A*_541_ values for GBS-treated wells to the *A*_541_ values for the wells with blood incubated with water.

### GBS BspC and capsule flow cytometry

Flow cytometry to determine BspC and capsule expression was performed as described in (80). Briefly, bacterial stocks were washed in sterile PBS containing 0.5% bovine serum albumin (BSA) (VWR) then incubated with a purified monoclonal anti-serotype III antibody or a purified monoclonal anti-serotype Ia isotype control at a 1:10,000 dilution, washed via centrifugation, and labeled with a donkey anti-mouse IgM conjugated to AlexaFluor647 (Invitrogen) at a 1:2,000 dilution. All incubations were performed at 4°C with shaking. Samples were washed again then resuspended and read on a FACScalibur flow cytometer (BD Biosciences), and analyzed using FlowJo (v10) software. The monoclonal antibodies were provided by John Kearney at the University of Alabama at Birmingham.

To stain for surface BspC expression, GBS were grown to OD_600_ of 0.25 in EndoGRO-MV culture medium (Millipore) in order to mimic host infection conditions, pelleted by centrifugation, resuspended in PBS and frozen *L. lactis* strain stocks were thawed and washed in buffer. Approximately, 1 x 10^6^ CFU of each strain was incubated with either adsorbed anti-BspC antibody or adsorbed anti-rabbit IgG at a 1:50 dilution at 4° C, overnight, with rotation. The next day, bacteria were washed via centrifugation, and labeled with a donkey anti-rabbit IgG conjugated to AlexaFluor488 (Invitrogen) at a 1:2,000 dilution for 45 minutes at room temperature with rotation. Samples were washed again then resuspended and read on a FACScalibur flow cytometer (BD Biosciences), and analyzed using FlowJo (v10) software.

### Immunofluorescent staining of GBS

Bacteria were grown to an OD_600_ of 0.25 in EndoGRO-MV culture medium (Millipore), the bacteria suspension was smeared on charged glass slides (Fisher), and the slides were fixed with 4% paraformaldehyde for 30 min at room temperature. The slides were blocked with 3% BSA for 1 hour, then incubated with rabbit antibodies to BspC or IgG at a 1:50 dilution followed by donkey anti-rabbit conjugated to AlexaFluor488 (Invitrogen). Bacteria were imaged using a BZ-X710 fluorescent microscope (Keyence).

### Scanning electron microscopy

Bacteria were grown to log phase and were then fixed for 10 min using a one-step method with 2.5% glutaraldehyde, 1% osmium tetraoxide, 0.1M sodium cacodylate. Bacteria were collected on 0.4 μM polycarbonate filters by passing the solution through a swinnex device outfitted on a 10 mL syringe. The filters were dehydrated through a series of increasing ethanol concentrations and then dried in a Tousimis SAMRI-790 critical point drying machine. The dried filters were mounted on SEM sample stubs with double-sided carbon tape, coated with 6nm platinum using a Quorom Q150ts high-resolution coater and imaged with a FEI FEG450 scanning electron microscope.

### Cell lines and infection assays

Cells of the well-characterized human cerebral microvascular endothelial cell line (hCMEC/D3), referred to here as hCMEC were obtained from Millipore and were maintained in an EndoGRO-MV complete medium kit supplemented with 1 ng/ml fibroblast growth factor-2 (FGF-2; Millipore) (81–84). Hela57A were provided by Marijke Keestra-Gounder at the University of Colorado, Anschutz Medical Campus and cultured in DMEM (Corning Cellgro) containing 10% fetal bovine serum (Atlanta Biologicals). HEK293T cells were obtained from Origene and cultured in DMEM containing 10% fetal bovine serum and 2mM L-glutamine (Thermo Fisher). The lentiviral expression plasmid pLenti-C-Myc-DDK harboring the human vimentin gene (NM_003380, pLenti-VIM) was obtained from Origene. To generate lentiviruses, HEK293T cells were transfected with the pLenti-VIM plasmid in combinations with the packaging plasmid psPAX2 and the envelope plasmid pMD2.G (Addgene) using TransIT_293 transfection reagent (Mirus). After an 18 h incubation, the culture supernatant containing lentiviruses was harvested and filtered through a 0.45 μm syringe filter to remove cellular debris. The viral titer was 10^6^ to 10^7^ transduction units (TU) per mL. 10^5^ fresh HEK293T cells were infected with lentiviruses at a MOI of 5 for 24 h in the presence of 10 μg/mL polybrene (Sigma). The empty lentiviral expression plasmid pLenti-mock was used as a vector control. Assays to determine the total number of cell surface-adherent or intracellular bacteria were performed as describe previously (10). Briefly, bacteria were grown to mid-log phase to infect cell monolayers (1 × 10^5^ CFU, at a multiplicity of infection [MOI] of 1). Total cell-associated GBS and *L. lactis* were recovered following a 30 min incubation, while intracellular GBS were recovered after 2 h infection and 2 h incubation with 100 μg gentamicin (Sigma) and 5 μg penicillin (Sigma) to kill all extracellular bacteria. Cells were detached with 0.1 ml of 0.25% trypsin-EDTA solution and lysed with addition of 0.4 ml of 0.025% Triton X-100 by vigorous pipetting. The lysates were then serially diluted and plated on THB agar to enumerate bacterial CFU. For antibody pre-treatment assays, hCMEC were incubated with 0.3μg/ml antibodies for 30 min prior to infection with GBS. The mouse monoclonal antibody to vimentin, clone V9 (Abcam), the rabbit polyclonal antibody to N-terminal vimentin (Sigma), and the isotype controls (VWR) were used. Total cell-associated GBS were recovered following a 1h incubation. For luciferase assays, Hela57A cells were infected with 1 x 10^6^ CFU (MOI, 10) GBS for 90 min. Cells were then lysed and luciferase activity quantified using a luciferase assay system (Promega) according to manufacturer’s instructions.

### Mouse model of hematogenous GBS meningitis

We utilized a mouse GBS infection model as described previously (10, 26, 27). Briefly, 8-week old male CD-1 mice (Charles River), 129S WT, or 129S-*Vim^tm1Cba^*/MesDmarkJ (Vimentin KO) (Jackson Laboratory) were injected intravenously with 1 × 10^9^ CFU of wild-type GBS or the isogenic Δ*bspC* mutant for a high dose challenge, or 1 × 10^8^ CFU for a low dose challenge. At the experimental endpoint mice were euthanized and blood, lung, and brain tissue were collected. The tissue was homogenized, and the brain homogenates and lung homogenates as well as blood were plated on THB agar for enumeration of bacterial CFU.

### Histology

Mouse brain tissue was frozen in OCT compound (Sakura) and sectioned using a CM1950 cryostat (Leica). Sections were stained using hematoxylin and eosin (Sigma) and images were taken using a BZ-X710 microscope (Keyence).

### Brain flow cytometry

At 48 h post-infection with 1 × 10^8^ CFU of GBS, mice were euthanized then perfused to replace blood with PBS. The entire mouse brain was harvested from each animal and the tissue was processed with the Multi-Tissue Dissociation kit #1 following the Adult brain dissociation protocol (Miltenyi Biotec). Cells were resuspended in MACS buffer (Miltenyi Biotec) and incubated with antibodies to Ly6C conjugated to BV421, CD45 conjugated to PE, CD11b conjugated to FITC, and Ly6G conjugated to APC (Invitrogen) at 1:200 dilution, UltraLeaf anti-mouse CD16/CD32 Fc block (Biolegend) at 1:400 dilution, and fixable viability dye conjugated to eFLuor506 (eBioscience) at 1:1000 dilution for 1 h, then fixed (eBioscience). Cells were counted using a Countess automated cell counter (Invitrogen), read on a Fortessa X-20 flow cytometer (BD Biosciences), and analyzed using FlowJo (v10) software. Gates were drawn according to fluorescence minus one (FMO) controls.

### RT-qPCR and ELISA

GBS were grown to mid-log phase and 1 × 10^6^ CFU (MOI, 10) were added to hCMEC monolayers and incubated at 37°C with 5% CO_2_ for 4 h. Cell supernatants were collected, the cells were then lysed, total RNA was extracted (Machery-Nagel), and cDNA was synthesized (Quanta Biosciences) according to the manufacturers’ instructions. Primers and primer efficiencies for IL-8, CXCL-1, and GAPDH (glyceraldehyde-3-phosphate dehydrogenase) were utilized as previously described (85). IL-8 and CXCL-1 from hCMEC supernatants, and KC and IL-1β from mouse brain homogenates were detected by enzyme-linked immunosorbent assay according to the manufacturer’s instructions (R&D systems).

### Immunofluorescent staining of hCMEC and HEK293T cells

hCMEC were grown to confluency on collagenized coverslips (Fisher). Following a 1 h infection, cells were washed with PBS to remove non-adherent bacteria and fixed with 4% paraformaldehyde (Sigma) for 30 min. For Figure 4, cells were incubated with 1% BSA in PBS with 0.01% Tween-20 (Research Products International) to block non-specific binding for 15 min, then incubated with a rabbit antibody to p65 (Sigma) at a 1:200 dilution overnight at 4°C. Coverslips were then washed with PBS and incubated with donkey anti-rabbit conjugated to Cy3 (Jackson Immunoresearch) at a 1:500 dilution for 1 h at room temperature. For Figure 6, cells were incubated with 1% BSA in PBS for 15 min, then with antibodies to vimentin (Abcam) and GBS (Genetex) at a 1:200 dilution overnight at 4°C. Following washes with PBS and an incubation with donkey anti-mouse conjugated to Cy3 and donkey anti-rabbit conjugated to 488 secondary antibodies (ThermoFisher) at a 1:500 dilution for 1 h at room temperature, coverslips were washed with PBS and mounted onto glass microscopy slides (Fisher) with VECTASHIELD mounting medium containing DAPI (Vector Labs). Cells were imaged using a BZ-X710 fluorescent microscope (Keyence). Quantification of GBS and vimentin co-localization was performed by counting the number of GBS that co-localized with vimentin and dividing by the total number of GBS in each field. For figure 7, HEK293T cells were incubated with the antibody to vimentin followed by a FITC-conjugated secondary antibody. Cells were imaged using a Cytation 5 fluorescent microplate reader (BioTek).

### 2-dimensional electrophoresis (2-DE), Far Western blot, and mass spectrometry analysis

Membrane proteins of hCMEC cells were enriched using a FOCUS membrane protein kit (G Biosciences, St. Louis, MO), dissolved in rehydration buffer (7M urea, 2M thiolurea, 1% TBP, and 0.2% ampholytes 3-10 NL), and quantified using 2D Quant kit (GE Healthcare, Piscataway, NJ). Proteins (100μg) were loaded on 7-cm long immobilized pH gradient (IPG) strips with non-linear (NL) 3-10 pH gradient (GE Healthcare). Isoelectric focusing was carried out in Multiphor II electrophoresis system (GE Healthcare) in three running phase (phase 1: 250V/0.01h, phase 2: 3500V/1.5h, and phase 3: 3500V/ 4.5h). The second dimension SDS-PAGE was carried out using 12.5% acrylamide gels in duplicate. One gel was stained with Coomassie Blue G250 (Bio-Rad, Hercules, CA) for mass spectrometry analysis. The other gel was transferred to a PVDF membrane for far Western blot analysis.

The PVDF membrane was denaturated and renaturated as described in (86), followed by incubation in a blocking solution (5% skim milk in PBS) for 1 h. Recombinant BspC was biotinylated using a EZ-Link Sulfo-NHS-Biotin kit (ThermoFisher Scientific, Waltham, MA). The PVDF membrane was probed with the biotinylated BspC (100µg) in a blocking solution overnight at 4°C. After washing three times with a washing buffer (PBS, 0.05% Tween-20), the PVDF membrane was incubated with an antibody conjugated to streptavidin-horse radish peroxidase (HRP). Interacting proteins were detected by adding enhanced chemiluminescence (ECL) reagents (ThermoFisher Scientific) and visualized by x-ray film exposure. The protein spots from far Western blot were aligned to the corresponding protein spots in the Coomassie stained gel. The identified spots were excised and digested in gel with trypsin (Worthington, Lakewood, NJ). Peptide mass spectra were collected on MALDI-TOF/TOF, (ABI 4700, AB Systems, Foster City, CA) and protein identification was performed using the automated result dependent analysis (RDA) of ABI GPS Explorer softwareV3.5. Spectra were analyzed by the Mascot search engine using the Swiss protein database.

### Bacterial two-hybrid assay

A bacterial adenylate cyclase two-hybrid assay was performed as in (32) and following manufacturer’s instructions (Euromedex). Briefly, plasmids containing T25-Vimentin and BspC-T18 were transformed into *E. coli* lacking *cyaA* and *E. coli* were plated on LB plates containing X-gal (Sigma). To measure β -galactosidase activity, Miller assays were performed according to standard protocols (87). Briefly, *E. coli* were grown in 0.5mM IPTG (Sigma), then permeabilized with 0.1% SDS (Sigma) and toluene (Sigma). ONPG (Research Products International) was added and absorbance was measured at 600, 550, and 420nm.

### Microscale thermophoresis

Three independent MST experiments were performed with His-tagged BspC labelled using the Monolith His-Tag Labeling Kit RED-tris-NTA 2^nd^ Generation (NanoTemper Technologies) according to manufacturer’s instructions. The concentration of labelled BspC was kept constant at 10nM. Vimentin was purchased from Novus Biologicals and titrated in 1:1 dilutions to obtain a series of 16 titrations ranging in concentration from 20 μM to 0 μM. Measurements were performed in standard capillaries with a Monolith NT.115 Pico system at 20% excitation power and 40% MST power (NanoTemper Technologies).

### Imaging flow cytometry

Following a 1 h infection with GBS, hCMEC were washed with PBS to remove non-adherent GBS. Cells were collected using a cell scraper (VWR) and resuspended in PBS containing 10% FBS and 1% sodium azide (Sigma). Cells were incubated with primary antibodies to vimentin (Abcam) and GBS (Genetex) at 1:500 dilution for 1 h at 4°C followed by donkey anti-mouse conjugated to Cy3 and donkey anti-rabbit conjugated to 488 (ThermoFisher) secondary antibodies. Cells were then fixed (eBioscience) and analyzed using an ImageStream X imaging flow cytometer (Amnis).

### Data analysis

GraphPad Prism version 7.0 was used for statistical analysis and statistical significance was accepted at *P* values of <0.05. (*, *P* < 0.05; **, *P* < 0.005; ***, *P* < 0.0005; ****, *P* < 0.00005). Specific tests are indicated in figure legends.

## FIGURE LEGENDS

Figure S1. (A-C) Flow cytometry using a polyclonal rabbit antibody to BspC to show expression of BspC in WT COH1 (A), Δ*bspC* mutant (B), and the complemented (C) GBS strains. (D-G) Immunofluorescent staining of WT COH1 (D), Δ*bspC* mutant (E), and the complemented (F) GBS strains using the BspC antibody to show surface localization of BspC protein. (G) Negative staining control. Scale bar is 5 μm. (H) Growth curves for WT GBS and the Δ*bspC* mutant in THB. (I) Hemolysis assay comparing hemolysis of sheep blood by WT GBS and the Δ*bspC* mutant. Representative data of one of at least three independent experiments are shown. Error bars represent the standard deviation of mean in one experiment. Data were analyzed using an unpaired t test. (J and K) Flow cytometry using a monoclonal antibody to the serotype III capsule to determine the presence of capsule in WT GBS (J) and the Δ*bspC* mutant (K) and a monoclonal antibody to the serotype Ia capsule as an isotype control. (L) Quantification of capsule flow cytometry data shown in (J) and (K).

Figure S2. Flow cytometry to show BspC surface expression in *L. lactis* containing the pMSP empty plasmid (A) and *L. lactis* containing the pMSP.*bspC* vector (B)

Figure S3. (A) Kaplan-Meier plot showing survival of mice challenged with either WT 515 GBS or the isogenic Δ*bspC* mutant. (B-D) Tissue bacterial counts for mice infected with WT 515 and 90356 GBS and the isogenic Δ*bspC* mutants. 48h post-infection, mice were sacrificed and bacterial loads in brain (B), lung (C), and blood (D) were quantified. Statistical analysis: (A) Log-rank test. (B-D) Two-way ANOVA with Sidak’s multiple comparisons test. *, *P* < 0.0005;**, *P* < 0.005.

Figure S4. (A) Far Western blot analysis of hBMEC membrane proteins using biotinylated BspC protein. Two spots (I and II) were identified on the x-ray film and aligned to the Coomassie stained gel. (B) Electrospray ionization-tandem mass spectrometry identifies spots I and II as vimentin. (C) The amino acid sequence of human vimentin, with the peptide sequences identified in the MS analysis underscored and bolded. (D) Control Far Western blot with the streptavidin antibody conjugated to HRP only. (E) Representative MST dose response curve quantifying the dissociation constant for the interaction between BspC and vimentin.

Figure S5. Immunofluorescent staining of WT 129 (A) and 129 *Vim*^-/-^ (B) brain tissue sections with an antibody to vimentin and with tomato lectin to label blood vessels. Nuclei were labelled with DAPI. Scale bar is 50 μm.

Figure S6. (A) Kaplan-Meier plot showing survival of WT 129 mice or 129 *Vim*^-/-^ mice challenged with GBS Δ*bspC* mutant. (B) 48h post-infection, mice were sacrificed and bacterial loads in brain, lung, and blood were quantified.

## ACKNOWLEDGEMENTS

We thank Dr. Marijke Keestra-Gounder (University of Colorado, Anschutz Medical Campus) for the Hela57A reporter cell line, John Kearney (University of Alabama at Birmingham) for GBS monoclonal antibodies, Tibor Pechan of the Mississippi State University Institute for Genomics, Biocomputing & Biotechnology (IGBB) for assistance with the mass spectrometry, Dr. Felipe Almeida de Pinho-Ribeiro at Harvard Medical School for assistance with brain cell flow cytometry analysis, Jeremy Rakhola at the Rocky Mountain Regional VA Medical Center (RMRVAMC) Flow Core, Shaun Bevers at the University of Colorado Anschutz Medical Campus Biophysics Core, and Catherine Back (University of Bristol, UK) for provision of purified *S gordonii* CshA protein. Also to Katilynn Croom and Jeffrey Kavanaugh for technical support. This study was supported by the Rees Stealy Research Foundation/ SDSU Heart Institute and San Diego Chapter ARCS Scholarships to L.D., the NIH 5T32AI007405-28 to B.L.S., the Coordenação de Aperfeiçoamento de Pessoal de Nível Superior - Brasil (CAPES) - Finance Code 001 to G.F.S., the NIH DE016690 to H.F.J., and the NIH/NINDS R01NS051247 to K.S.D.

