## Supplementary figures and images for "The Group B Streptococcal surface antigen I/II protein, BspC, interacts with host vimentin to promote adherence to brain endothelium and inflammation during the pathogenesis of meningitis"

### Supplemental Figure 1

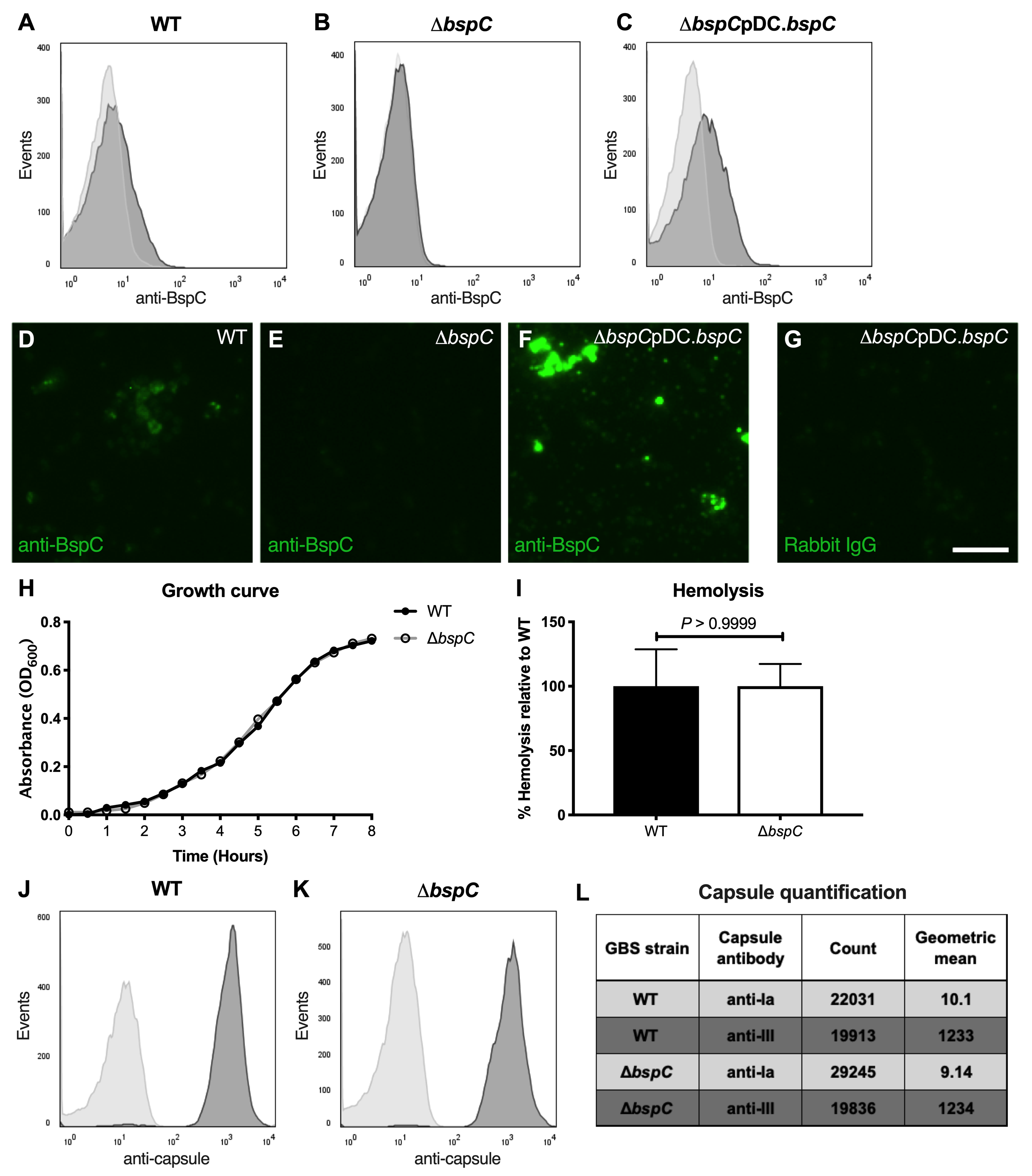

### Supplemental Figure 2

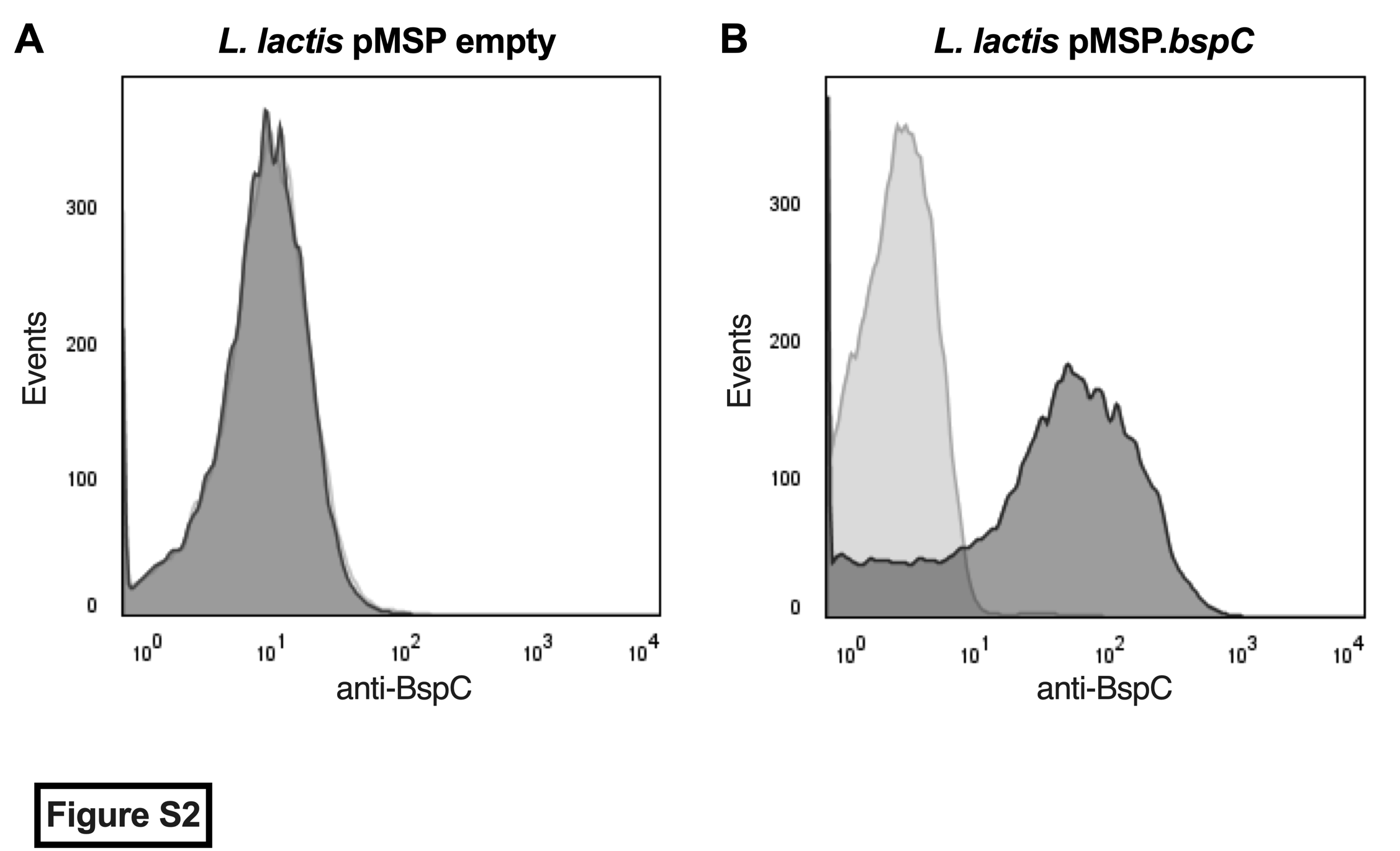

### Supplemental Figure 3

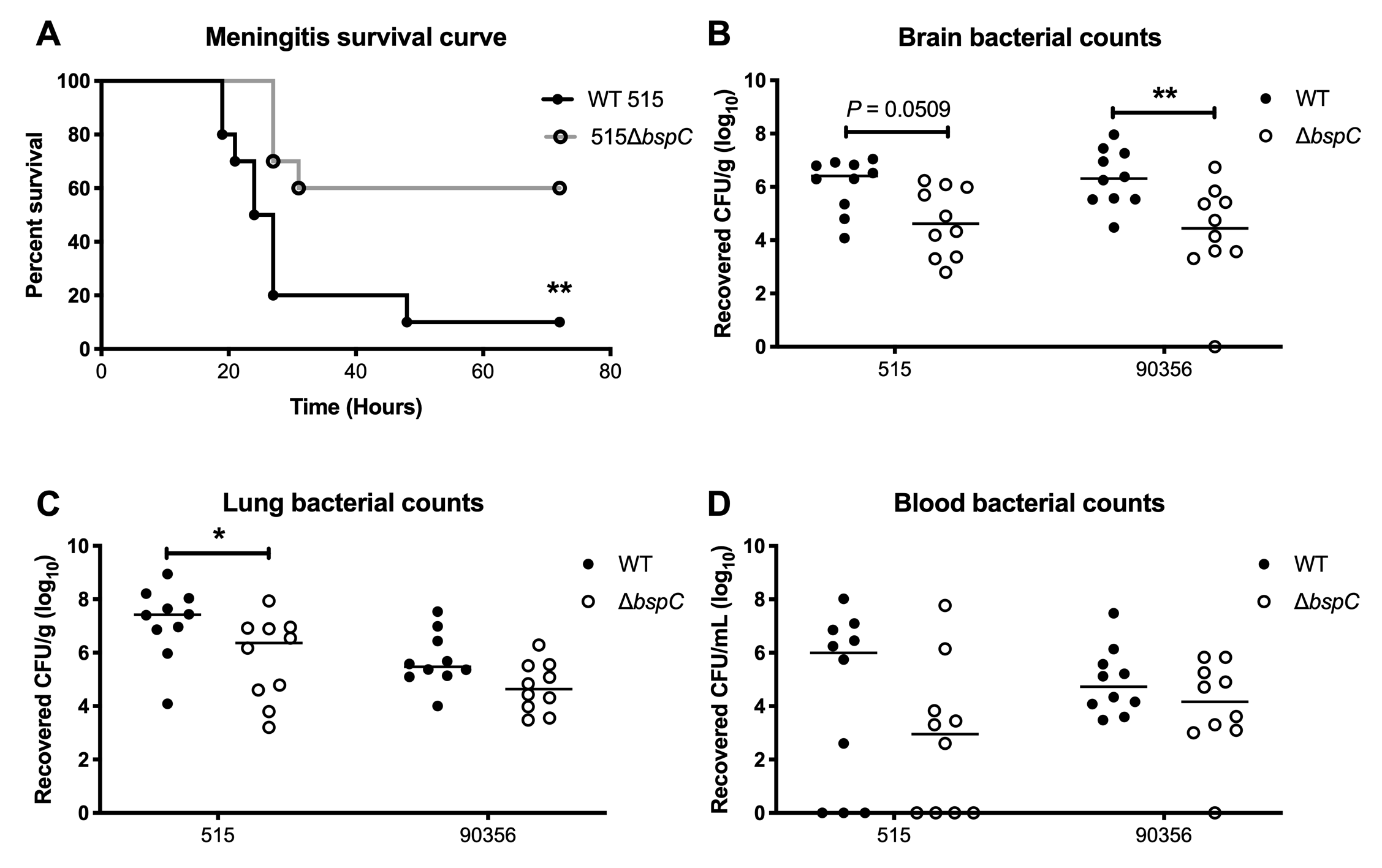

### Supplemental Figure 4

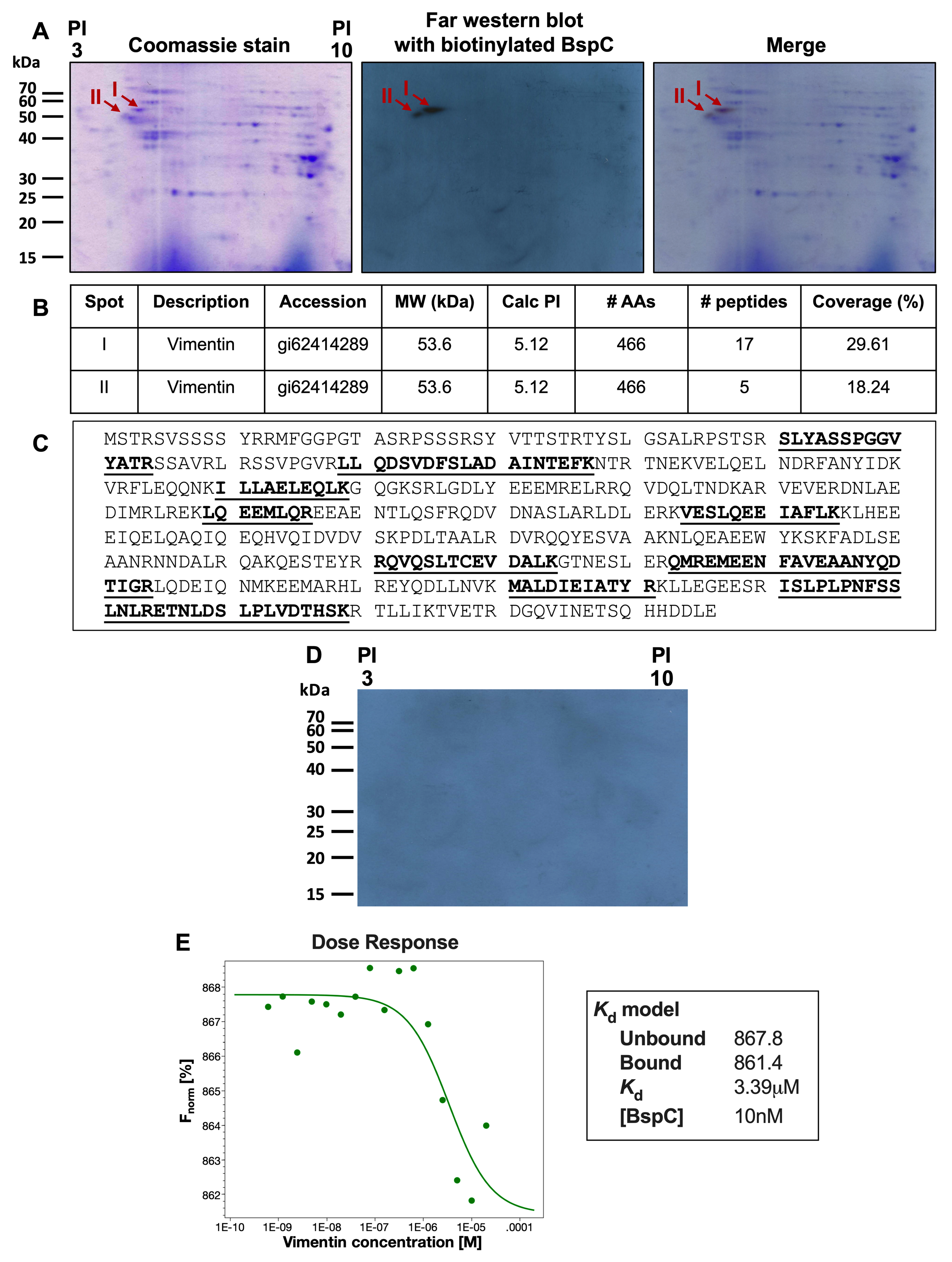

### Supplemental Figure 5

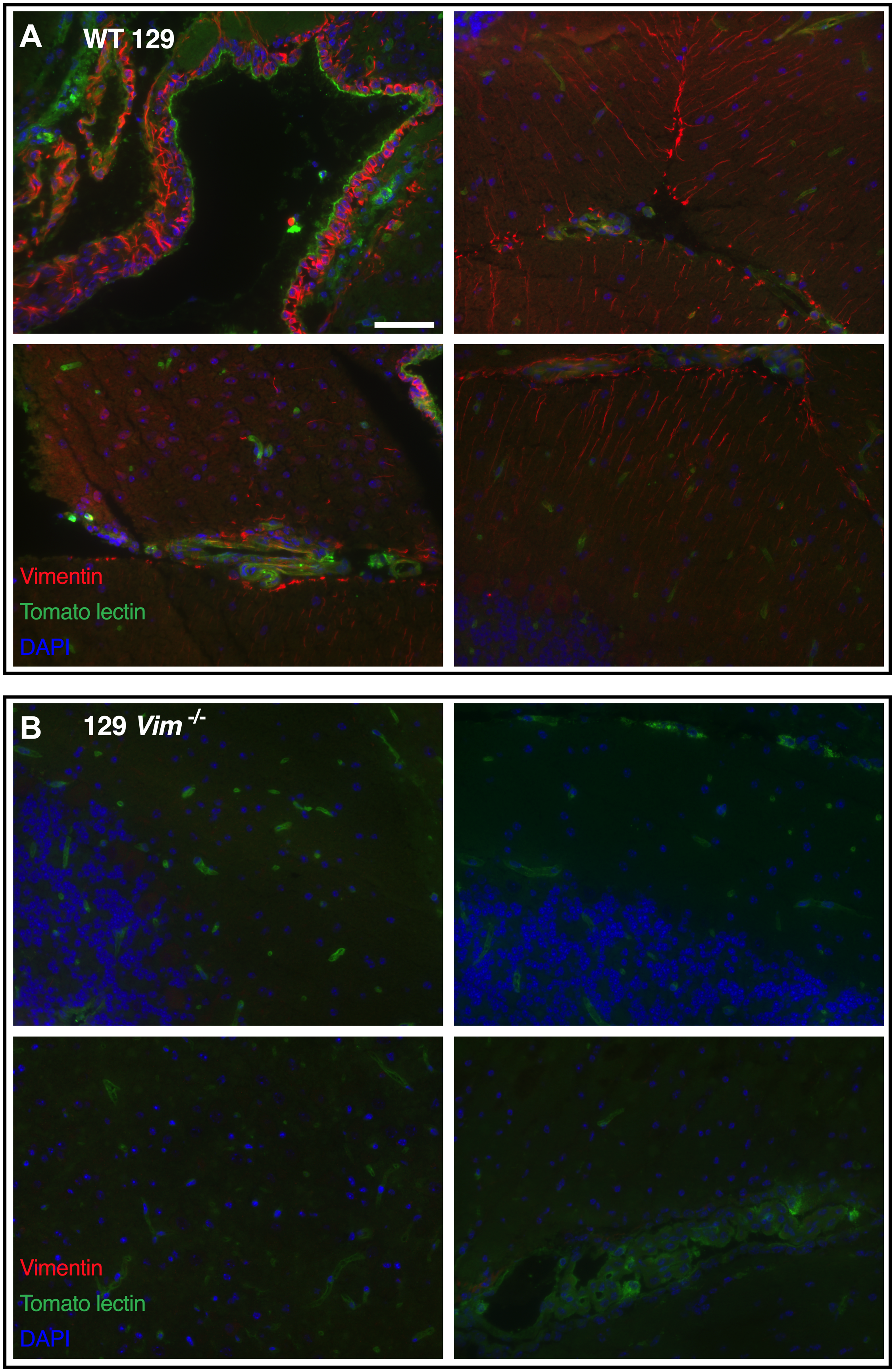

### Supplemental Figure 6

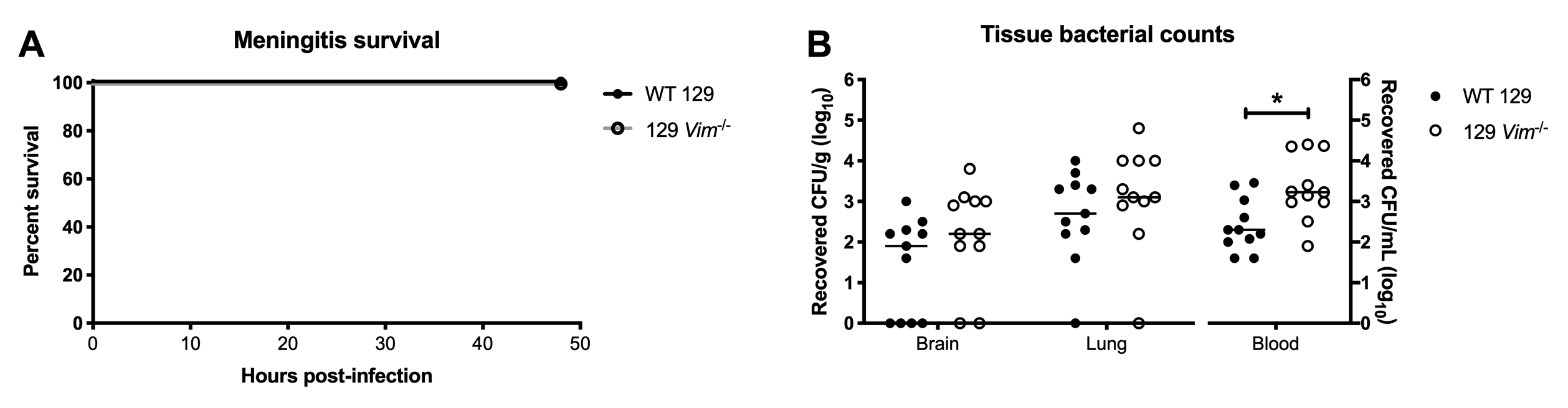
